# 14-3-3 mitigates alpha-synuclein aggregation and toxicity in the *in vivo* preformed fibril model

**DOI:** 10.1101/2020.11.17.387415

**Authors:** Rachel Underwood, Mary Gannon, Aneesh Pathak, Navya Kapa, Sidhanth Chandra, Alyssa Klop, Talene A. Yacoubian

**Author notes:** **Corresponding Author: Talene A. Yacoubian, MD, PhD, 1719 Sixth Avenue South, Civitan International Research Center, Room 510A, Birmingham, AL 35294,**.

## Abstract

Alpha-synuclein (αsyn) is the key component of proteinaceous aggregates termed Lewy Bodies (LBs) that pathologically define a group of disorders known as synucleinopathies, including Parkinson’s Disease (PD) and Dementia with Lewy Bodies (DLB). αSyn is hypothesized to misfold and spread throughout the brain in a prion-like fashion. Transmission of αsyn necessitates the release of misfolded αsyn from one cell and the uptake of that αsyn by another, in which it can template the misfolding of endogenous αsyn upon cell internalization. 14-3-3 proteins are a family of highly expressed brain proteins that are neuroprotective in multiple PD models. We have previously shown that 14-3-3θ acts as a chaperone to reduce αsyn aggregation, cell-to-cell transmission, and neurotoxicity in the *in vitro* pre-formed fibril (PFF) model. In this study, we expanded our studies to test the impact of 14-3-3s on αsyn toxicity in the *in vivo* αsyn PFF model. We used both transgenic expression models and adenovirus associated virus (AAV)-mediated expression to examine whether 14-3-3 manipulation impacts behavioral deficits, αsyn aggregation, and neuronal loss in the PFF model. 14-3-3θ transgene overexpression in cortical and amygdala regions rescued social dominance deficits induced by PFFs at 6 months post injection, whereas 14-3-3 inhibition by transgene expression of the competitive 14-3-3 peptide inhibitor difopein in the cortex and amygdala accelerated social dominance deficits. The behavioral rescue by 14-3-3θ overexpression was associated with delayed αsyn aggregation induced by PFFs in these brain regions. Conversely, 14-3-3 inhibition by difopein in the cortex and amygdala accelerated αsyn aggregation and cortical pyramidal neuron loss induced by PFFs. 14-3-3θ overexpression by AAV in the substantia nigra (SN) also delayed αsyn aggregation in the SN and partially rescued PFF-induced dopaminergic cell loss in the SN. 14-3-3 inhibition in the SN accelerated nigral αsyn aggregation and increased PFF-induced dopaminergic cell loss. These data indicate a neuroprotective role for 14-3-3θ against αsyn toxicity *in vivo*.

## Introduction

Alpha-synuclein (αsyn) is a critical protein whose aggregation and transmission from cell to cell has been implicated in the neurodegenerative process in Parkinson’s disease (PD) and Dementia with Lewy Bodies (DLB). αSyn is a highly expressed brain protein whose endogenous function is not well understood but likely includes regulation of synaptic transmission [5, 8, 23]. Thorough understanding of the mechanisms that regulate the aggregation, transmission, and toxicity of this protein could lead to new targets for therapeutic intervention in these disorders. We recently observed that the chaperone-like protein 14-3-3θ is a critical regulator of the release, oligomerization, and toxicity of αsyn in several cellular models [46]. 14-3-3θ is a member of the highly homologous 14-3-3 protein family, which are multifunctional proteins that play a role in protein folding, protein trafficking, neurite growth, and cell survival among other cellular roles [13, 16, 22, 31, 33, 42, 50]. 14-3-3θ is found to be colocalized with αsyn in Lewy Bodies in both PD and DLB [4, 14]. We have observed a reduction in 14-3-3θ expression in transgenic αsyn mice and in soluble 14-3-3 levels in DLB brains [6, 20, 48, 49]. Additionally, aberrant phosphorylation of 14-3-3θ has been noted in both PD and DLB brains [20]. 14-3-3θ overexpression protects against both neurotoxin and mutant LRRK2 toxicity, while 14-3-3 inhibition increases toxicity [7, 16, 36, 49].

Given 14-3-3s’ roles in protein folding and trafficking, we recently examined the impact of 14-3-3s on αsyn cell-to-cell transmission and toxicity in two separate cellular models: the paracrine αsyn model and the *in vitro* αsyn fibril model [46]. We found that 14-3-3θ reduces αsyn transfer and toxicity by inhibiting αsyn oligomerization, seeding, and internalization, whereas 14-3-3 inhibition accelerates the αsyn seeding and cell-to-cell transmission in these cellular models [46]. In the study described here, we expanded our work to examine the impact of 14-3-3θ on αsyn toxicity *in vivo* using the αsyn preformed fibril (PFF) model. Here we describe the effect of 14-3-3θ overexpression or 14-3-3 inhibition on behavioral deficits, αsyn inclusion formation, and neuronal loss in the PFF model. We observed that 14-3-3θ overexpression reduced social dominance deficits, delayed αsyn inclusion formation, and reduced neuronal loss, while pan 14-3-3 inhibition with the peptide inhibitor difopein accelerated behavioral deficits, αsyn inclusion formation, and neuronal loss in the PFF model.

## Methods

### Mice

Mice were used in accordance with the guidelines of the National Institute of Health (NIH) and University of Alabama at Birmingham (UAB) Institutional Animal Care and Use Committee (IACUC). All animal work performed in this study was approved by UAB’s IACUC. Transgenic mice expressing human 14-3-3θ under the neuronal promoter Thy1.2 were previously developed by our group [16, 46]. Transgenic mice expressing difopein-enhanced yellow fluorescent protein (eYFP) under the neuronal promoter Thy1.2 were obtained from Dr. Yi Zhou at Florida State University[29]. 14-3-3θ and difopein hemizygous mice were crossed with C57BL/6J mice from The Jackson Laboratory (catalog #000664; RRID:IMSR_JAX:000664) to produce transgenic mice for stereotactic injections. C57BL/6J mice from The Jackson Laboratory (catalog #000664; RRID:IMSR_JAX:000664) were purchased and homozygous bred to produce wildtype (WT) mice for AAV experiments.

### Fibril preparation

Recombinant full-length mouse αsyn protein was prepared as previously described and generously supplied by Dr. Laura Volpicelli-Daley and Dr. Andrew West [10, 43]. Before stereotaxic injection in mice, fibrils were generated by incubating purified mouse monomeric αsyn at a concentration of 5 mg/ml in 50mM Tris (pH 7.4) with 166 mM KCl with constant agitation at 700 rpm at 37ºC for 7 days. Immediately prior to injection, αsyn fibrils were sonicated with a water bath sonicator (QSonica, Newton CT) for 1 hour at A=30 at 4ºC. Sonicated αsyn fibrils were analyzed by dynamic light scattering (DLS) on a DynaPro NanoStar (Wyatt Technology, Santa Barbara CA) to ensure sonicated fibril average radius was between 10-100 nm prior to injection.

### AAV preparation and injection

AAV2/CBA-IRES2-eGFP-WPRE (AAV-GFP) and AAV2/CBA-1433thetaV5his-IRES-eGFP-WPRE (AAV-14-3-3θ) were constructed as previously described [7, 38]. Male and female C57BL/6 mice from Jackson Laboratories were deeply anesthetized with 5% isoflurane and maintained at 0.25%-4% during surgery for stereotactic injection at 8 weeks with AAV into the SN [anteroposterior (AP): −3.0 from bregma; mediolateral (ML):-1.3 from midline; dorsoventral (DV): −4.6 below dura]. Mice were injected with 2 μl of either AAV-GFP (titer: 1.8E+12 vg/ml, viral genomes/ml) or AAV-14-3-3θ (titer: 7.0E+11 vg/ml) at a rate of 0.25 μl/min using a microinjection pump.

### Stereotactic injections of PFFs

Mice at 8-12 weeks of age were deeply anesthetized with 5% isoflurane and maintained at 0.25%-4% during surgery for PFF injection. WT mice injected with AAV-GFP or AAV-14-3-3θ/GFP at 8 weeks of age were then injected at 12 weeks of age with PFFs. 14-3-3θ, difopein, or WT littermates were injected at 8-12 weeks of age with PFFs. Mice were unilaterally injected with 5 μg of αsyn fibrils or monomer (2 μl of 2.5 mg/ml) at a flow rate of 0.250 μl/minute into the dorsolateral striatum (AP: 0.2 mm from bregma; ML: −2 mm from midline; DV: −2.6 mm below dura) according to a previously established protocol [19]. Post-surgery mice were allowed a minimum of 15 minutes recovery time on a heating pad. All mice were observed until fully awake and non-drowsy to ensure successful recovery. Mice were given buprenorphine (1mg/kg) 20 minutes prior to surgery and the day after to minimize pain and discomfort.

### Immunohistochemistry

Mice were perfused with PBS followed by 4% paraformaldehyde using a forced pump system. After dissection, brains were sliced by microtome in 40 μm thick coronal sections. Every sixth section from the anterior brain (including the sensorimotor and striatal regions) was stained for pS129 αsyn or NECAB1 immunohistochemistry, while every fourth section from the posterior sections (including the substantia nigra) was stained for pS129 αsyn or TH immunohistochemistry. For pS129 αsyn and TH immunohistochemistry, sections were quenched with 0.6% hydrogen peroxide in methanol followed by antigen retrieval (10 mM sodium citrate, 0.05% Tween-20, pH 6.0) for 1 hour at 37° C. Sections were then blocked in 5% normal goat serum (NGS) with 0.3% Triton X-100 and incubated for 48 hours in pS129-αsyn antibody (Abcam #51253) or 24 hours in tyrosine hydroxylase antibody (Pelfreez #40101) in 1.5% NGS. After washing, sections were incubated with goat anti-rabbit IgG biotinylated secondary antibody (Vector Laboratories #BA-1000) for 4 hours at 4°C. After washing with TBS, sections were incubated in ABC solution (Vector) for 30 minutes and developed using ImmPACT DAB Chromagen Solution (Vector). Following washing with TBS, sections were mounted on slides and progressively dehydrated with ethanol and Histo-clear. Slides were cover-slipped with Permount (Avantar) and imaged using an Olympus BX51 epifluorescence microscope. For pS129-αsyn positive inclusion counts, 3 sections in either the sensorimotor cortical regions (Bregma +1.54 mm to +0.62 mm) or the SN (Bregma −2.70 mm to −3.88 mm) per well were selected and quantitated using ImageJ with the rater blind to experimental conditions. One section was selected in the central and basolateral amygdala region (Bregma −0.82 mm to −1.34 mm) and quantitated using ImageJ with the rater blind to experimental conditions. αSyn inclusion counts were normalized per mm^2^ area.

For NECAB1 staining, sections were washed in TBS followed by antigen retrieval (10 mM sodium citrate, 0.05% Tween-20, pH 6.0) for 1 hour at 37° C. Sections were then blocked in 5% NGS with 0.1% Triton X-100 and incubated with anti-NECAB1 rabbit antibody (Sigma Millipore #HPA023629) and anti-NeuN (Thermofisher Scientific #14H6L24) mouse antibody overnight at 4°C. After washing with TBS, sections were incubated with Cy3-conjugated goat anti-mouse secondary antibody and Alexa488-conjugated goat anti-rabbit antibody for 2 hours at 4°C. Sections were mounted on slides, cover-slipped using ProLong Diamond Antifade mounting solution (Thermofisher Scientific), and imaged using an Olympus BX51 epifluorescence microscope. For NECAB1-positive neuronal counts, 3 sections in the sensorimotor cortical regions (Bregma +1.54 mm to +0.62 mm) per well were selected and quantitated using ImageJ with the rater blind to experimental conditions. NECAB1 counts were normalized to mm^2^ area.

### Stereology

Stereological estimates of TH-positive neuronal numbers were performed using the optical fractionator method of the StereoInvestigator 8.0 software from MBF Biosciences (Microbrightfield Inc., Williston, VT, USA), as previously described [7, 39]. For each animal, SNpc regions of every fourth section based on systematic random sampling were outlined according to published mouse atlas (Bregma −2.70 mm to −3.88 mm). A grid was placed randomly over the outlined region for counting. At each counting frame (50 μm × 50 μm) of the grid predetermined by the software setup, neurons with visible nuclei were counted within three-dimensional optical dissectors set to 20 μm with a 60x oil immersion objective using an Olympus BX51 Microscope. A 1-μm guard distance from the top and bottom of the section surface was excluded from each dissector. Section thickness was measured at every tenth counting frame on each section to obtain the actual thickness after tissue processing. The total number of neurons (Ntotal) was calculated using the equation: Ntotal = Ncounted × 1/ssf × 1/asf × 1/hsf, where Ncounted is the number of neurons counted, ssf is the section sampling fraction, asf is the area sampling fraction and hsf is the height sampling fraction. Coefficient of error (Gundersen, m = 1) was set to <0.1. Stereology estimates were done with the investigator blinded to the experimental condition.

### Behavior Tests

Behavior tests to assess motor and social functions were conducted 3 and 6 months after PFF injection. Mice were handled for 3 to 5 days before testing began and habituated to testing room for 30 minutes at the start of each testing day. Behavior tests began at least 1 hour after light/dark cycle switch and completed at least 1 hour before switching back to their dark cycle. In order to minimize stress to the animals, behavior tests were ordered from least to most stressful as follows: open field, wire hang, social dominance (tube test), pole test, and finally rotarod. Behavior was done primarily at 6 months post injection (mpi), but the difopein cortical cohort did undergo behavioral tests at 3 mpi. All behavior apparatuses were cleaned with 2% chlorohexidine between trials in order to minimize scent contamination between mice. Further, male mice were tested before female mice to minimize their exposure to female scent.

### Open Field

For open field, mice were placed with a 48 inch × 48 inch open arena with clear plexi-glass walls. Mice were videoed for 4 minutes using EthoVision software and analyzed for overall velocity and distance moved across all planes (vertical and horizontal).

### Wire Hang

A four limb wire hang test was performed, as previously described with some modifications [15]. An apparatus with a wire grid bottom and angled edges to prevent mice from crawling over was built by the UAB machine shop. This apparatus was attached to a ring stand approximately 1 meter high over a rat cage filled with bedding. The apparatus was inverted and a mouse was set on top. The apparatus was then flipped back over, so that the mouse was upside down and had to hang onto the wire grid in order to stay on. If the mouse was still hanging on after 60 seconds, it was removed and placed back in its home cage. A total of 2 trials per mouse were run, and all the mice in the cohort were run on the first trial before beginning the second trials, giving an interval of about 90 minutes between the two trials.

### Tube Test

For the tube test, mice were placed in a clear 12” long tube at opposing ends as previously described [3, 40]. Mice were not released until both mice had all four paws inside the tube. Male mice were tested in a 1½” diameter tube and females were tested in a 1” diameter tube to account for differences in size. Each PFF-injected WT mouse, monomer-injected transgenic 14-3-3θ or difopein mouse, and PFF-injected transgenic 14-3-3θ or difopein mouse was matched against 3 different monomer-injected WT mice in separate trials. Mice were given a 2-minute rest between rounds.

### Pole Test

At 3 mpi (aged 5-6 months), mice were placed facing upwards at the top of a ¼” diameter round 3-foot long pole. Each mouse was timed for the duration to turn downwards down the pole as well as descend to the bottom of the pole. At 6 mpi (mouse age 8-9 months), wire footholds were added to the pole to assist mice in descending safely due to age and weight gain.

### Rotarod

To further assess motor activity at 6 mpi, mice were placed on an accelerating rotarod apparatus with speed increasing from 5 to 35 rpm over 60 seconds. Mice were given 3 days of training on the apparatus and 2 days of testing. All mice underwent 3 trials per day with an inter-trial rest time of 2 minutes on training days and 5 minutes on assessment days. No mice failed to fall during the allotted 60s.

### Statistical Analysis

GraphPad Prism 8 (La Jolla, CA) was used for statistical analysis of experiments. Data were analyzed by either Student’s t-test, one-way ANOVA, or two-way ANOVA, followed by post-hoc pairwise comparisons using Tukey’s multiple comparison tests. Statistical significance was set at p ≤ 0.05. All the details of experiments can be found in the results section or figure legends. All data values are presented as mean ± SEM. ANOVA related statistics (F statistic, p values) are noted in the results section, while the post-hoc test results are found in the figure legends. For t-tests, the t statistic and p values are noted in the results section.

## Results

To test the impact of 14-3-3 proteins on αsyn pathogenesis *in vivo,* we examined the effect of 14-3-3θ overexpression or 14-3-3 inhibition on behavioral deficits, αsyn inclusion formation, and neuronal loss in the *in vivo* fibril model (Fig. 1). We previously created a transgenic mouse that overexpresses 14-3-3θ tagged with the HA tag under the Thy1.2 promoter [16, 46]. This transgenic mouse expresses HA-tagged 14-3-3θ in neurons located in the cortex, hippocampus, amygdala, and other areas, but no HA-tagged 14-3-3θ is detected in dopaminergic neurons in the SN (Supp. Fig. 3b). This mouse was used to examine the impact of 14-3-3θ manipulation in the cortex and amygdala (Fig. 1a). We also examined the impact of 14-3-3θ overexpression in the SN by using an adeno-associated virus (AAV) expressing 14-3-3θ-GFP that was injected into the SN by stereotactic means (Fig. 1b). For testing the impact of 14-3-3 inhibition with the pan-14-3-3 peptide inhibitor difopein, we used 2 different lines expressing difopein-eYFP under the Thy1.2 promoter [29]: 1) line 132, which primarily expresses difopein-eYFP in neurons in the cortex and amygdala but not within the substantia nigra, and 2) line 166, which expresses difopein-eYFP in tyrosine hydroxylase-positive neurons in the SN but does not have expression in the cortex (Fig. 5a, d; Supp. Fig. 4a).

**Figure 1.**
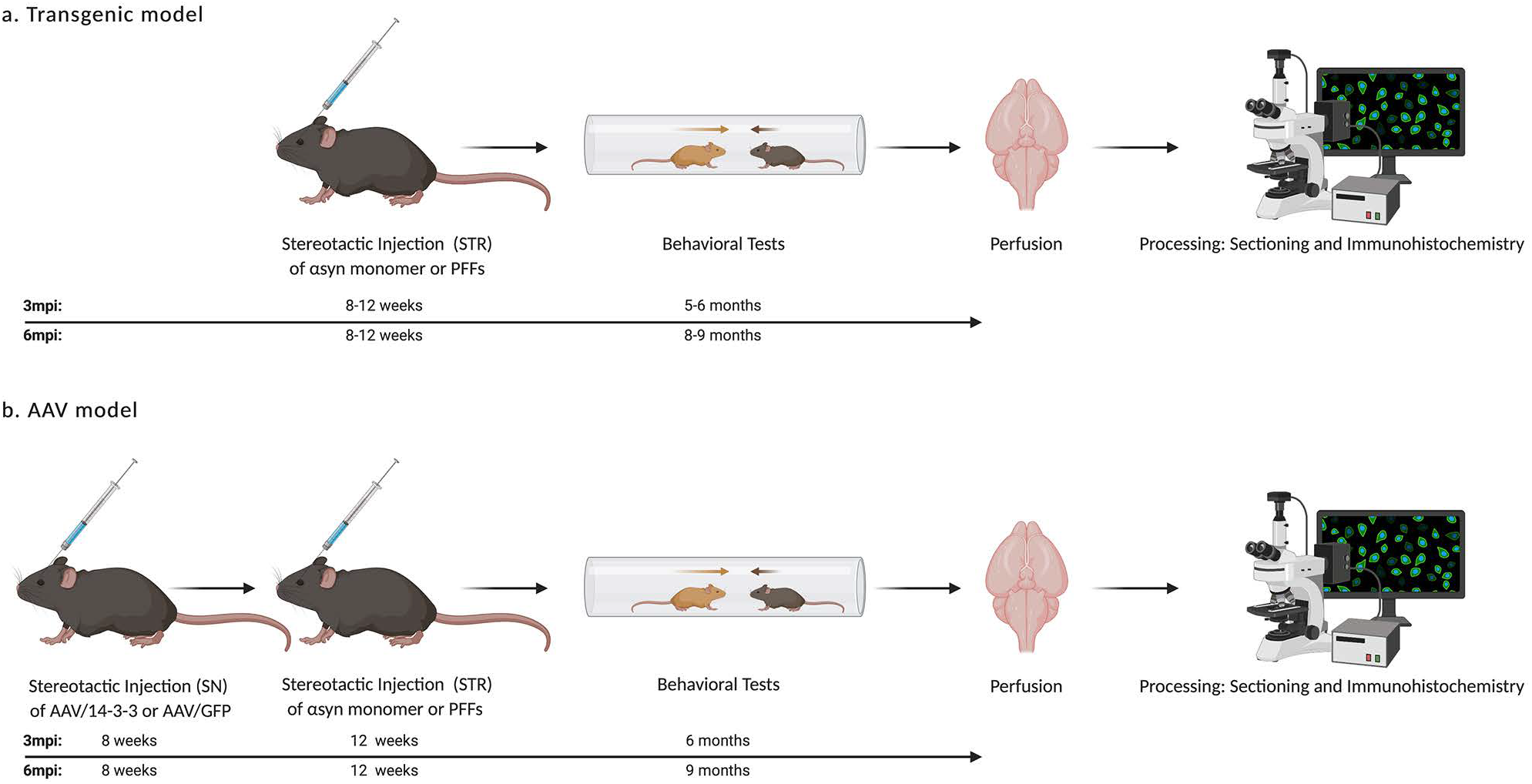
Experimental design of *in vivo* PFF experiments. a) Transgenic difopein-eYFP expressing or 14-3-3θ-overexpressing transgenic mice and matching WT littermates were given a unilateral stereotactic injection of mouse αsyn monomer or PFFs in the striatum (STR) at 8-12 weeks of age. At 3 or 6 months post injection (mpi), mice underwent behavioral testing. Mice were then perfused, and their brains were sectioned by microtome and processed for immunohistochemical staining. b) WT mice were given a unilateral injection of AAV-14-3-3θ/GFP or AAV-GFP in the substantia nigra (SN) at 8 weeks of age. 4 weeks after AAV inoculation, mice were given an ipsilateral stereotactic injection of αsyn monomer or PFFs in the STR at 12 weeks of age. Mice underwent behavioral testing at 6 mpi. Mice were then perfused, and their brains were sectioned by microtome and processed for immunohistochemical staining.

### 14-3-3θ overexpression reduces behavioral deficits induced by αsyn fibrils

Previous studies have shown motor and/or cognitive behavioral effects in response to αsyn PFF injection into the dorsolateral striatum [12, 18, 40]. WT and 14-3-3θ transgenic littermates were injected unilaterally with monomeric or fibrillary αsyn (5 μg) into the dorsolateral striatum at 8 to 12 weeks of age. Six months after PFF injection, we examined motor and non-motor behaviors in these mice. While some studies have shown a motor deficit in mice injected with PFFs at 6 mpi [18], we did not observe a consistent motor deficit in WT animals injected with PFFs, as determined by the pole test or rotarod test at 6 mpi (Supp. Fig. 1b, c). We also did not observe any differences in monomeric or fibrillar αsyn-injected 14-3-3θ mice with regard to motor function on the pole test (2-way ANOVA: genotype F (1, 44) = 2.217e-005, p=0.9963; PFF treatment F (1, 44) = 0.01661, p=0.8980; interaction F (1, 44) = 0.08812, p=0.7680; Supp. Fig. 1b) or rotarod test (2-way ANOVA: genotype F (1, 45) = 0.08034, p=0.7781; PFF treatment F (1, 45) = 2.507, p=0.1203; interaction F (1, 45) = 0.02684; p=0.8706; Supp. Fig. 1c). Similarly, no difference was observed in velocity (2-way ANOVA: genotype F (1, 45) = 0.2994, p = 0.5870; PFF treatment F (1, 45) = 1.079, p = 0.3044; interaction F (1, 45) = 0.008179, p = 0.9283) or distance traveled (2-way ANOVA: genotype F (1, 45) = 0.2940, p = 0.5903; PFF treatment F (1, 45) = 1.088, p = 0.3024; interaction F (1, 45) = 0.007373, p = 0.9320) between WT and 14-3-3θ transgenic mice, whether injected with monomeric or fibrillar αsyn, in the open field test (Supp. Fig. 1a). We did observe a strong effect of PFF injections in WT animals in the social dominance test, which is thought to be a measure of prefrontal cortical and amygdala function [3, 21, 37, 45, 51], at 6 mpi (Fig. 2a). PFF-injected transgenic 14-3-3θ mice with 14-3-3θ overexpression primarily in the cortex showed a rescue of the social dominance deficit observed in WT littermates injected with PFFs (1-way ANOVA: F (2, 59) = 5.581; p = 0.0060; Fig. 2a).

**Figure 2.**
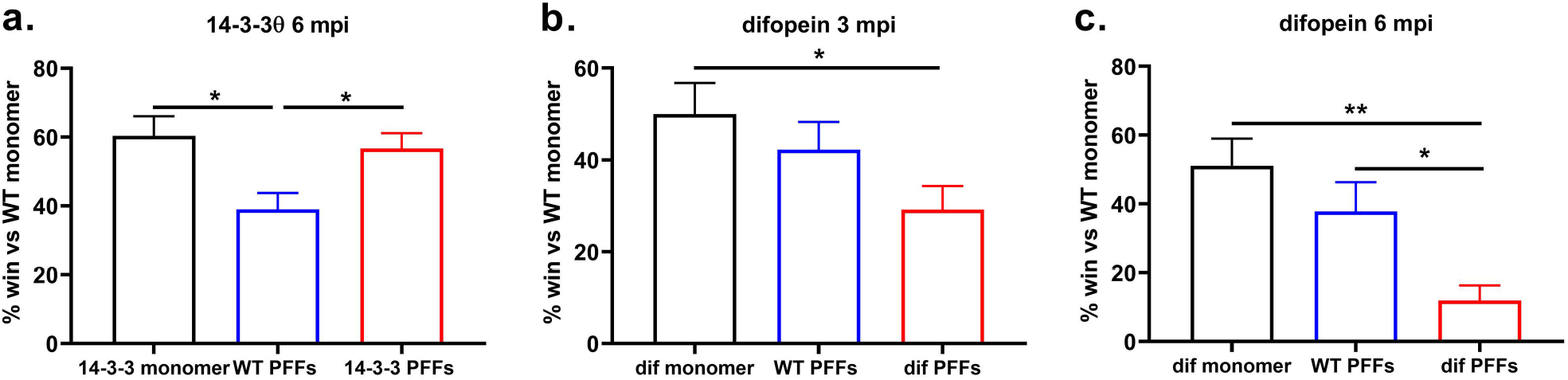
14-3-3θ overexpression reduces and 14-3-3 inhibition increases social dominance deficits induced by fibrillar αsyn. a) 14-3-3θ overexpression increased average win rate in comparison to WT PFF-injected mice. Quantification of win rate of 14-3-3θ mice injected with monomer, 14-3-3θ mice injected with PFFs, or WT mice injected with PFFs in the tube test matched against monomer-injected WT mice at 6 mpi. Each mouse was matched against 3 separate WT mice injected with monomer over 3 individual rounds, and the win rate per mouse was determined across these 3 trials. n=17-25 mice per group. *p<0.05 (Tukey’s multiple comparison test). Error bars represent SEM. b) Difopein transgenic expression in the cortex and amygdala decreased average win rate in comparison to PFF-injected WT mice. Quantification of difopein mice injected with monomer, difopein mice injected with PFFs, and WT mice injected with PFFs at 3 mpi in the tube test matched against monomer-injected WT mice. Mice were evaluated over 3 individual rounds. n=14-16 per group. *p<0.05 (Tukey’s multiple comparison test). Error bars represent SEM. c) Quantification of difopein mice injected with monomer, difopein mice injected with PFFs, and WT mice injected with PFFs at 6 mpi in the tube test matched against monomer-injected WT mice. Mice were evaluated over 3 individual rounds. n=14-15 per group. *p<0.05, **p<0.01 (Tukey’s multiple comparison test). Error bars represent SEM.

Since our transgenic 14-3-3θ line does not demonstrate 14-3-3θ overexpression in the nigra, we tested the impact of 14-3-3θ overexpression in the SN using an adeno-associated virus (AAV) expressing 14-3-3θ-GFP [7]. WT mice were stereotactically injected with AAV-GFP or AAV-14-3-3θ/GFP into the SN at 8 weeks of age, and then monomeric or fibrillar αsyn was injected into the ipsilateral dorsolateral striatum four weeks later. No significant motor deficit was observed in mice injected with AAV-GFP or AAV-14-3-3θ/GFP with or without PFFs in the wire hang test at 6 mpi (2-way ANOVA: genotype F (1, 52) = 1.554, p = 0.2181; PFF treatment F (1, 52) = 2.342, p = 0.1320; interaction F (1, 52) = 0.2655, p = 0.6086; Supp. Fig. 2a). While there was a slight PFF effect on the pole test (2 way ANOVA: genotype F (1, 52) = 0.02336, p = 0.8791; PFF treatment F (1, 52) = 4.251, p=0.0442; interaction F (1, 52) = 0.7770, p = 0.3821) and on the open field test (2 way ANOVA: genotype F (1, 52) = 0.04865, p = 0.8263; PFF treatment F (1, 52) = 10.01, p = 0.0026, interaction (1, 52) = 0.2352, p = 0.6297), no significant differences were observed between GFP mice injected with monomer vs. GFP mice injected with PFFs or between GFP mice injected with PFFs and 14-3-3θ mice injected with PFFs on either pole test or open field testing at 6 mpi (Supp. Fig. 2a).

### 14-3-3 inhibition exacerbates behavioral deficits induced by αsyn fibrils

We next examined whether inhibition of 14-3-3s with the pan-14-3-3 peptide inhibitor difopein affects behavioral deficits in the *in vivo* PFF model. Transgenic mice expressing difopein in the cortex showed a deficit in social dominance after PFF injection that was not observed in WT mice at 3 mpi (1-way ANOVA: F (2, 42) = 3.139, p = 0.05; Fig. 2b). At 6 mpi, the win rate on the social dominance test was lower in difopein mice injected with PFFs compared to WT mice injected with PFFs (1-way ANOVA: F (2, 41) = 7.359, p = 0.0019; Fig. 2c). PFF treatment showed an overall slight increase in velocity (2-way ANOVA: genotype F (1, 56) = 0.4789 p = 0.4918; PFF effect F (1, 56) = 10.57, p = 0.0020; interaction F (1, 56) = 0.2741, p = 0.6027) and distance traveled (2-way ANOVA: genotype F (1, 56) = 0.1289, p = 0.7209; PFF effect F (1, 56) = 11.94, p = 0.0011; interaction F (1, 56) = 0.6675, p = 0.4174) in mice on the open field test, but no dramatic differences were noted between individual experimental groups at 6 mpi (Supp. Fig. 1d).

Similar to that observed with the AAV-14-3-3θ injections into the SN, expression of difopein in the SN did not impact motor function significantly. WT and nigral difopein mice did not demonstrate any motor deficit on the rotarod test at 6 mpi (2-way ANOVA: genotype F (1, 56) = 0.6888, p = 0.4101; PFF treatment F (1, 56) = 2.386, p = 0.1281; interaction F (1, 56) = 0.09658, p = 0.7571; Supp. Fig. 2b). PFF or genotype did not impact greatly distance traveled (2-way ANOVA: genotype effect F (1, 60) = 0.6861, p = 0.4108; PFF treatment F (1, 60) = 0.2927, p = 0.5905; interaction F (1, 60) = 5.109; p = 0.0274) or velocity (2-way ANOVA: genotype effect F (1, 60) = 0.2982, p = 0.5870; PFF treatment F (1, 60) = 0.03472, p = 0.8528; interaction F (1, 60) = 13.13, p = 0.0006) on the open field test at 6 mpi, although a slight interaction effect was noted (Supp. Fig. 2b).

### 14-3-3θ overexpression delays αsyn inclusion formation

Given the reversal of the social dominance deficit in 14-3-3θ mice, we next examined the impact of 14-3-3θ on αsyn inclusion formation. At 3 mpi, we observed a dramatic number of inclusions that stained positive for phosphorylated S129-αsyn (pS129-αsyn) in the sensorimotor cortex of WT mice injected with PFFs compared to WT mice injected with monomeric αsyn (Supp. Fig. 3a). By 6 mpi, inclusion numbers in WT mice were dramatically reduced (Fig. 3c, d). Other groups have also noticed a decline in αsyn inclusions over time in mice, presumably due to the loss of neurons that develop inclusions [1, 25, 27]. In 14-3-3θ transgenic mice injected with PFFs compared to WT mice injected with PFFs, we observed a 41% reduction in the number of inclusions that stained positive for pS129-αsyn in the cortex at 3 mpi (unpaired, two-tailed t-test: t_(13)_ = 2.754, p = 0.0164; Fig. 3b, c). However, at 6 mpi, 14-3-3θ mice injected with PFFs showed a 2.4-fold increase in inclusion counts in the cortex compared to WT mice injected with PFFs (unpaired, two-tailed t-test: t_(17)_ = 3.232, p = 0.0049; Fig. 3d, Supp. Fig. 3a). In the amygdala, we observed a 60% decrease in the number of pS129-αsyn positive inclusions at 3 mpi in 14-3-3θ transgenic mice injected with PFFs compared to WT mice injected with PFFs (unpaired, two-tailed t-test: t_(15)_ = 2.757, p = 0.0147) but then a 60% increase in the number of inclusions in the amygdala at 6 mpi (unpaired, two-tailed t-test: t_(13)_ = 2.193, p=0.0471) in 14-3-3θ transgenic mice injected with PFFs compared to WT mice injected with PFFs (Fig. 3b, c, d; Supp. Fig. 3a). No differences in inclusion counts were noted between 14-3-3θ and WT mice at either 3 mpi (unpaired, two-tailed t-test: t_(13)_ = 0.1161, p = 0.9094) or 6 mpi (unpaired, two-tailed t-test: t_(10)_ = 0.5746, p = 0.5782) in the SN, in which 14-3-3θ overexpression is not seen in these transgenic mice (Fig. 3b, c, d; Supp. Fig. 3). These findings suggest that inclusion formation in the cortex and amygdala is delayed by 14-3-3θ overexpression, and that higher levels of inclusions seen at 6 mpi in the 14-3-3θ mice compared to WT mice could be due to a reduction in neuronal loss.

**Figure 3.**
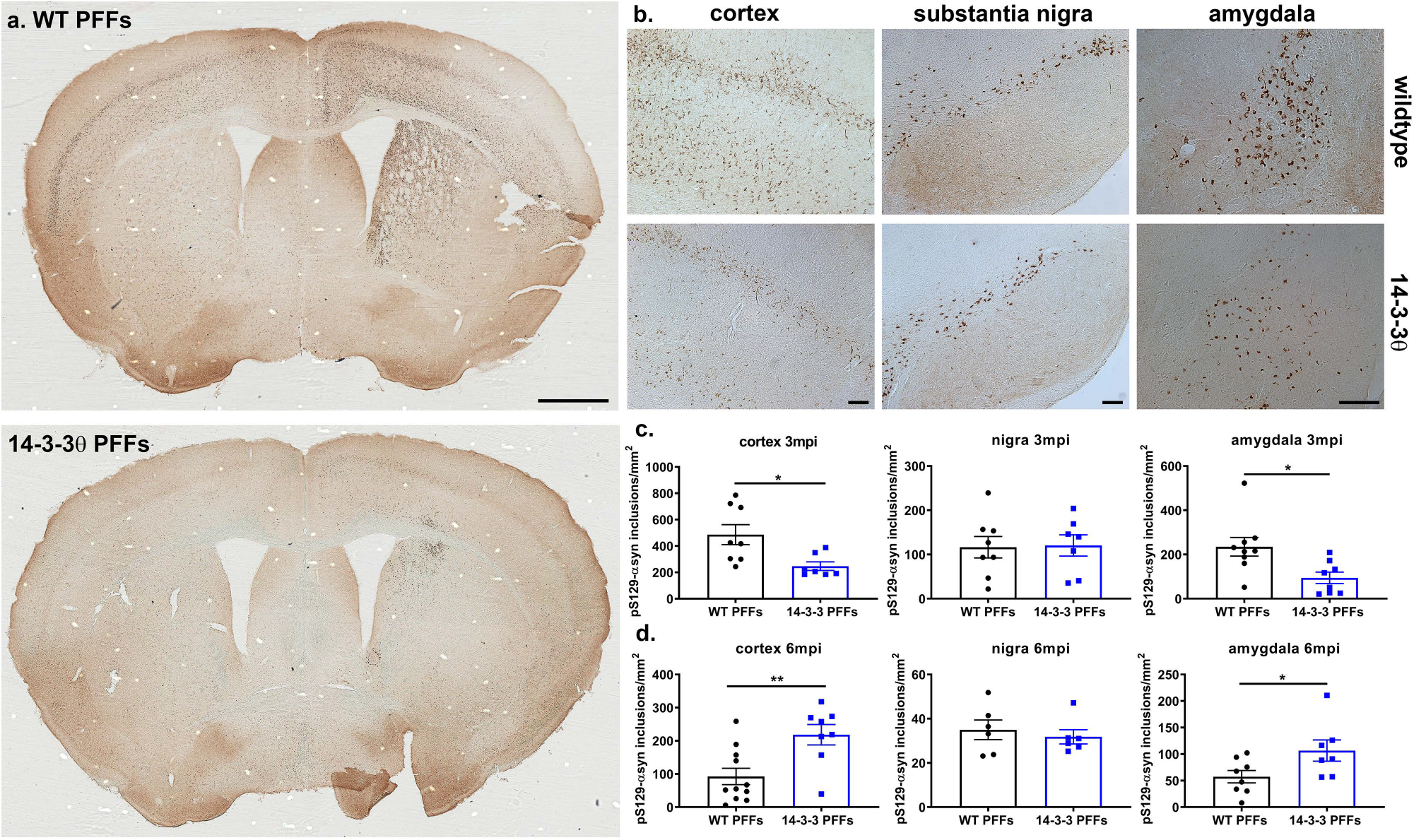
14-3-3θ overexpression delays αsyn inclusion formation in the cortex and amygdala. a) Representative images of PFF-injected WT and 14-3-3θ transgenic mice stained for pS129-αsyn demonstrate extensive αsyn inclusions in the STR and cortex in WT mice at 3 mpi. Scale bar = 1000 μm. b) Representative images of pS129-αsyn staining in PFF-injected WT and 14-3-3θ mice in the cortex, SN, and amygdala at 3 mpi. Scale bar = 100 μm. c) 14-3-3θ mice show decreased pS129-αsyn positive inclusion counts at 3 mpi in the cortex and amygdala, 2 areas with 14-3-3θ overexpression, but not the SN, which lacks 14-3-3θ overexpression. Quantification of pS129-αsyn positive inclusions in the cortex, SN, and amygdala of PFF-injected WT and 14-3-3θ mice at 3 mpi. n=7-9 per group. *p<0.05 (Student’s unpaired t-test). Error bars represent SEM. d) 14-3-3θ mice show increased pS129-αsyn positive inclusion counts in the cortex and amygdala but no change in the SN compared to WT mice at 6 mpi. Quantification of pS129-αsyn positive inclusions in the cortex, SN, and amygdala of PFF-injected WT and 14-3-3θ mice at 6 mpi. n=6-11 per group. *p<0.05, **p<0.01 (Student’s unpaired t-test). Error bars represent SEM.

Since our transgenic 14-3-3θ line does not demonstrate 14-3-3θ overexpression in the SN, we also examined nigral inclusion formation in the mice stereotactically injected with AAV-GFP or AAV-14-3-3θ/GFP in the SN (Fig. 4a). Consistent with our transgenic data, the number of inclusions positive for pS129-αsyn in the SN were decreased in mice injected with AAV-14-3-3θ/GFP compared to mice injected with AAV-GFP at 3 mpi (unpaired, two-tailed t-test: t_(25)_ = 2.229, p = 0.0350 Fig. 4b, d). By 6 mpi, the number of inclusions positive for pS129-αsyn in the SN were increased in mice injected with AAV-14-3-3θ/GFP compared to mice injected with AAV-GFP (unpaired, two-tailed t-test: t_(25)_ = 2.481, p = 0.0202; Fig. 4c, d). These findings suggest that inclusion formation in the nigra is also delayed by 14-3-3θ overexpression in the SN. The subsequent increase in inclusion formation at 6 mpi could reflect a reduction in neuronal loss in mice overexpressing 14-3-3θ in the SN.

**Figure 4.**
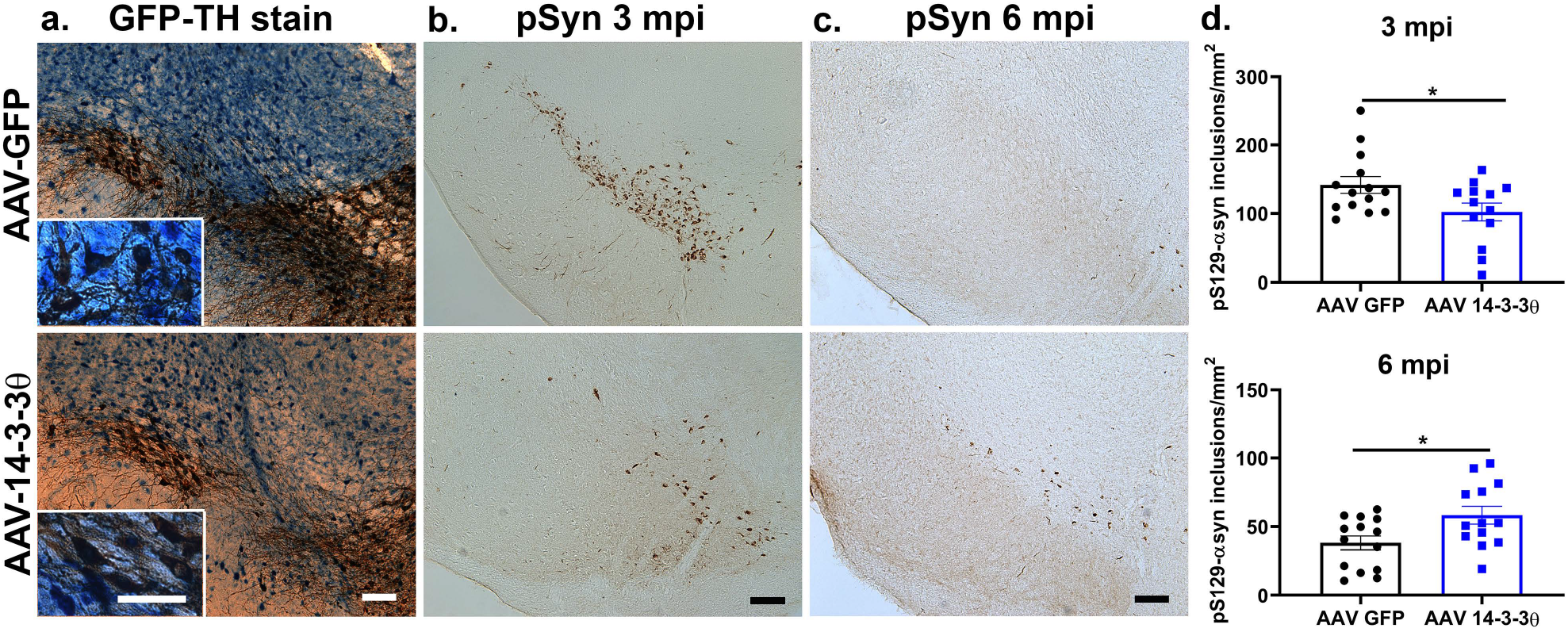
AAV-mediated 14-3-3θ overexpression delays αsyn inclusion formation in the substantia nigra. a) Representative images of immunohistochemistry for GFP (blue) and TH (brown) in brain sections containing the SN from AAV-GFP mice and AAV-14-3-3θ/GFP mice injected with αsyn monomers. Colocalization of GFP and TH staining indicates viral induction into dopaminergic nigral neurons in the targeted area. Scale bar = 100 μm for 10x image; scale bar = 50 μm for inset. b) Representative images of pS129-αsyn immunostaining in AAV-GFP mice and AAV-14-3-3θ/GFP mice injected with PFFs at 3 mpi. Scale bar = 100 μm. c) Representative images of pS129-αsyn immunostaining in AAV-GFP mice and AAV-14-3-3θ/GFP mice injected with PFFs at 6 mpi. Scale bar = 100 μm. d) AAV-mediated 14-3-3θ expression in PFF-injected mice decreases pS129-αsyn positive inclusion counts at 3 mpi and increases counts at 6 mpi in the SN. Quantification of nigral inclusions at 3 mpi and 6 mpi in AAV-GFP mice and AAV-14-3-3θ/GFP mice injected with PFFs. n=13-14 per group. *p<0.05 (Student’s t-test). Error bars represent SEM.

### 14-3-3 inhibition accelerates αsyn inclusion formation

We next examined whether inhibition of 14-3-3s with the pan-14-3-3 peptide inhibitor difopein affects αsyn inclusion formation in the *in vivo* PFF model. The difopein transgenic line that expresses difopein-eYFP in neurons in the cortex but not within the SN revealed increased inclusion counts in the cortex at 3 mpi compared to WT mice (unpaired, two-tailed t-test: t_(29)_ = 2.441, p = 0.0210; Fig. 5a, b, c). However, at 6 mpi, inclusion counts were significantly lower by 43% in the cortical difopein mice compared to WT injected with PFFs (unpaired, two-tailed t-test: t_(25)_ = 2.233, p = 0.0347; Fig. 5c). Similarly, in the amygdala, inclusion counts were increased by 31% at 3 mpi (unpaired, two-tailed t-test: t_(21)_ = 2.070, p = 0.05) but showed a non-significant decrease (53%) at 6 mpi (unpaired, two-tailed t-test: t_(15)_ = 2.021, p = 0.0616) in difopein mice compared to WT mice (Supp. Fig. 4). To test the impact of 14-3-3 inhibition on aggregation in nigral neurons, we measured inclusions in the nigral difopein transgenic line. Inclusion counts in the difopein nigral mice showed a non-significant increase at 3 mpi (unpaired, two-tailed t-test: t_(14)_ = 1.123, p = 0.2804) and decreased significantly at 6 mpi (unpaired, two-tailed t-test: t_(21)_ = 2.355, p = 0.0283) after PFF injection compared to WT mice (Fig. 5d, e, f). We conclude that 14-3-3 inhibition accelerates inclusion formation in response to αsyn fibrils, and that the reduction in inclusion counts at 6 mpi could reflect an increase in neuronal loss.

**Figure 5.**
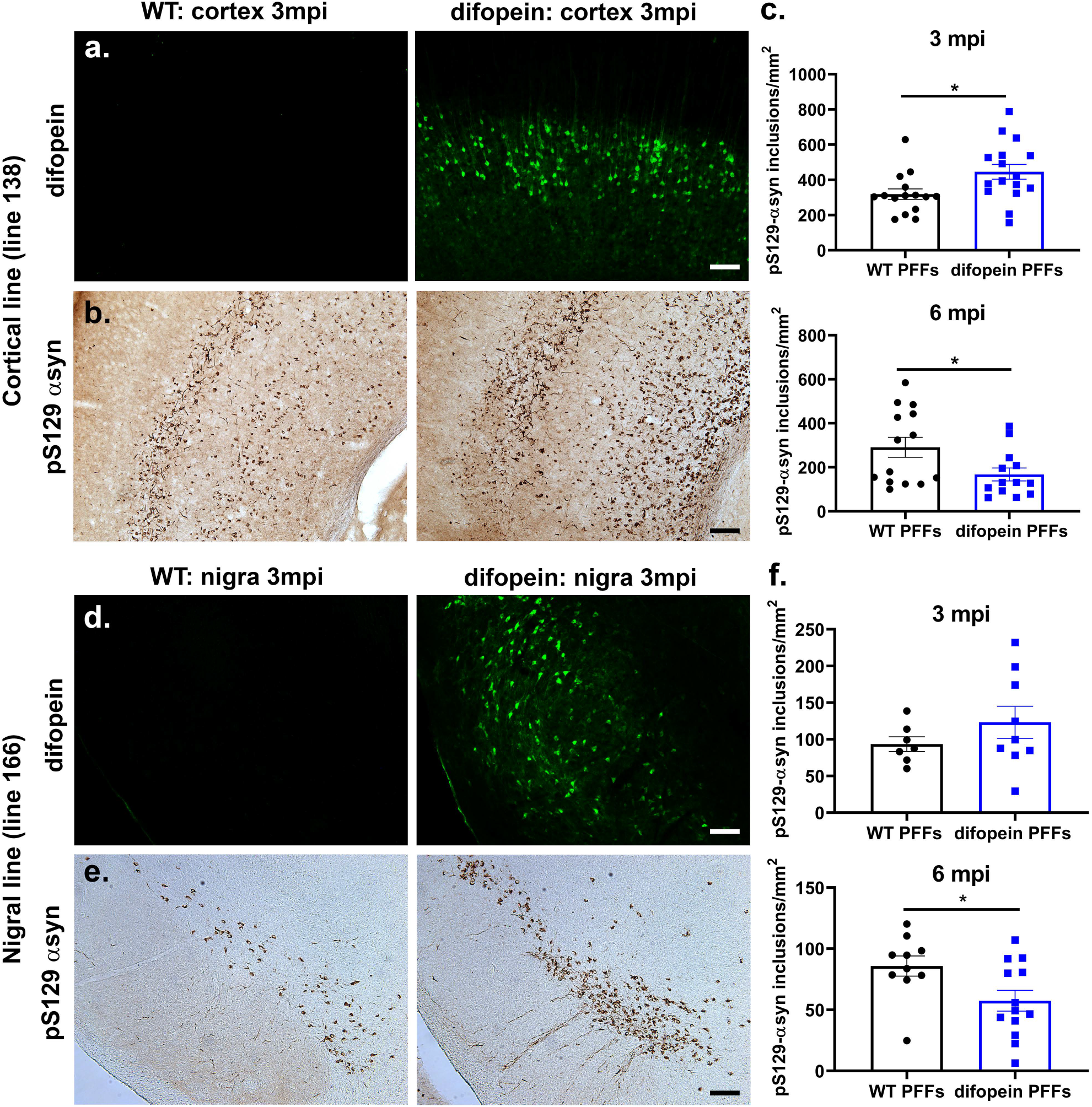
14-3-3 inhibition increases αsyn inclusion formation. a) Representative images of eYFP-difopein immunostaining in the cortex of WT and difopein (“cortical” line 138) mice at 3 mpi. GFP-difopein expression is found only in difopein mice. Scale bar = 100 μm. b) Representative images of pS129-αsyn immunostaining in the cortex of WT and difopein mice at 3 mpi. Scale bar = 100 μm. c) Difopein expression in the cortex increases inclusion counts at 3 mpi and decreases counts at 6 mpi in the cortex. Quantification of pS129-αsyn positive inclusions at 3 mpi and 6 mpi in the cortex of PFF-injected WT or difopein mice. n=15-16 per group at 3mpi; n=13-14 per group at 6mpi. *p<0.05 (Student’s t-test). Error bars represent SEM. d) Representative images of eYFP-difopein immunostaining in the SN of WT and difopein (“nigral” line 166) mice at 3 mpi. GFP-difopein expression is found only in difopein mice. Scale bar = 100 μm. e) Representative images of pS129-αsyn immunostaining in the SN of WT and difopein mice at 3 mpi. Scale bar = 100 μm. f) Difopein expression in in the nigra increases inclusion counts at 3 mpi and decreases counts at 6mpi in the SN. Quantification of pS129-αsyn positive inclusions at 3 mpi and 6 mpi in the SN of PFF-injected WT and difopein mice. n=7-9 per group at 3 mpi; n=10-13 per group at 6 mpi. *p<0.05 (Student’s t-test). Error bars represent SEM.

### 14-3-3s regulate neuronal loss induced by αsyn fibrils

As noted above, αsyn inclusions numbers are much higher at 3 months after PFF injection than at 6 months after injection in WT animals. This reduction in αsyn inclusion numbers over time is presumably secondary to the death of neurons that develop αsyn inclusions [25]. 14-3-3θ overexpression in either the nigra or cortex reduced αsyn inclusion numbers at 3 mpi, but we observed an increase in αsyn inclusions at 6 mpi in 14-3-3θ-overexpressing mice compared to control mice. We hypothesized that this delayed increase in inclusion formation with 14-3-3θ overexpression is due to the rescue of neurons that normally die in response to PFFs in control mice. To test the impact of 14-3-3θ overexpression on dopaminergic neuron loss in the SN, we performed stereological analysis of TH-positive neuronal counts in the nigra in mice injected with AAV-GFP or AAV-14-3-3θ/GFP. As expected, striatal PFFs induced a 25% loss in ipsilateral dopaminergic neuron counts in control mice injected with AAV-GFP into the ipsilateral nigra at 6 mpi (2-way ANOVA: genotype F (1, 43) = 0.8197, p = 0.3703; PFF treatment F (1, 43) = 10.01, p = 0.0029; interaction F (1, 43) = 2.514, p = 0.1202; Fig. 6a, b). In contrast, AAV-14-3-3θ/GFP mice injected with PFFs did not demonstrate a significant loss in dopaminergic neurons compared to AAV-14-3-3θ/GFP mice injected with monomeric αsyn at 6 mpi (Fig. 6a, b). This finding suggests that the increase in αsyn inclusions in 14-3-3θ-overexpressing mice at 6 mpi is likely due to a reduction in neuronal loss.

**Figure 6.**
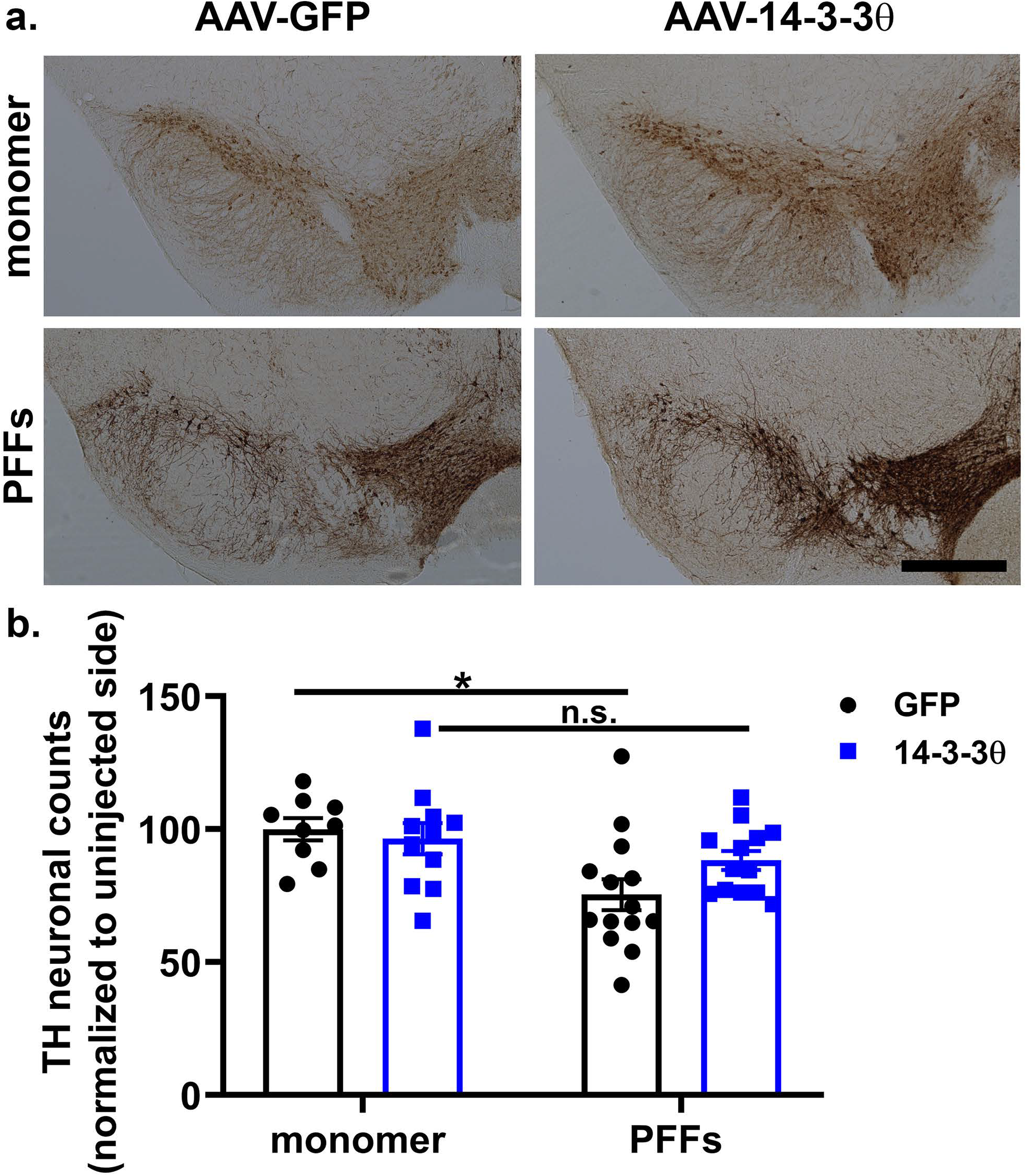
14-3-3θ overexpression mitigates nigral dopaminergic loss in response to PFFs. a) Representative images of TH immunohistochemistry in the SN of AAV-GFP and AAV-14-3-3θ/GFP mice injected with αsyn monomers or PFFs at 6 mpi. Scale bar = 500 μm. b) Stereological counts of TH-positive neurons in the SN of AAV-GFP and AAV-14-3-3θ/GFP mice injected with αsyn monomers or PFFs at 6 mpi. n=9-14 per group. *p<0.05 (Tukey’s multiple comparison test). n.s. = non-significant. Error bars represent SEM.

We next examined neuronal loss in response to PFFs in difopein-expressing mice. Stereological analysis of TH-positive counts in the nigra revealed a significant increase in ipsilateral dopaminergic neuronal loss in PFF-injected mice expressing difopein in the SN compared to WT mice after PFF injection at 6 mpi (2-way ANOVA: genotype F (1, 46) = 3.208, p = 0.0798; PFF treatment F (1, 46) = 58.27, p < 0.0001; interaction F (1, 46) = 4.309, p = 0.0435; Fig. 7a, b). Stereological analysis at 3 mpi showed a non-significant increase in dopaminergic neuron loss in difopein nigral mice, but no loss of dopaminergic neurons in WT mice injected with PFFs at 3 mpi (2-way ANOVA: genotype F (1, 26) = 3.282, p = 0.0816; PFF treatment F (1, 26) = 0.3335, p = 0.5686; interaction F (1, 26) = 1.105, p = 0.3028; Fig. 7b), as previously described by others [18].

**Figure 7.**
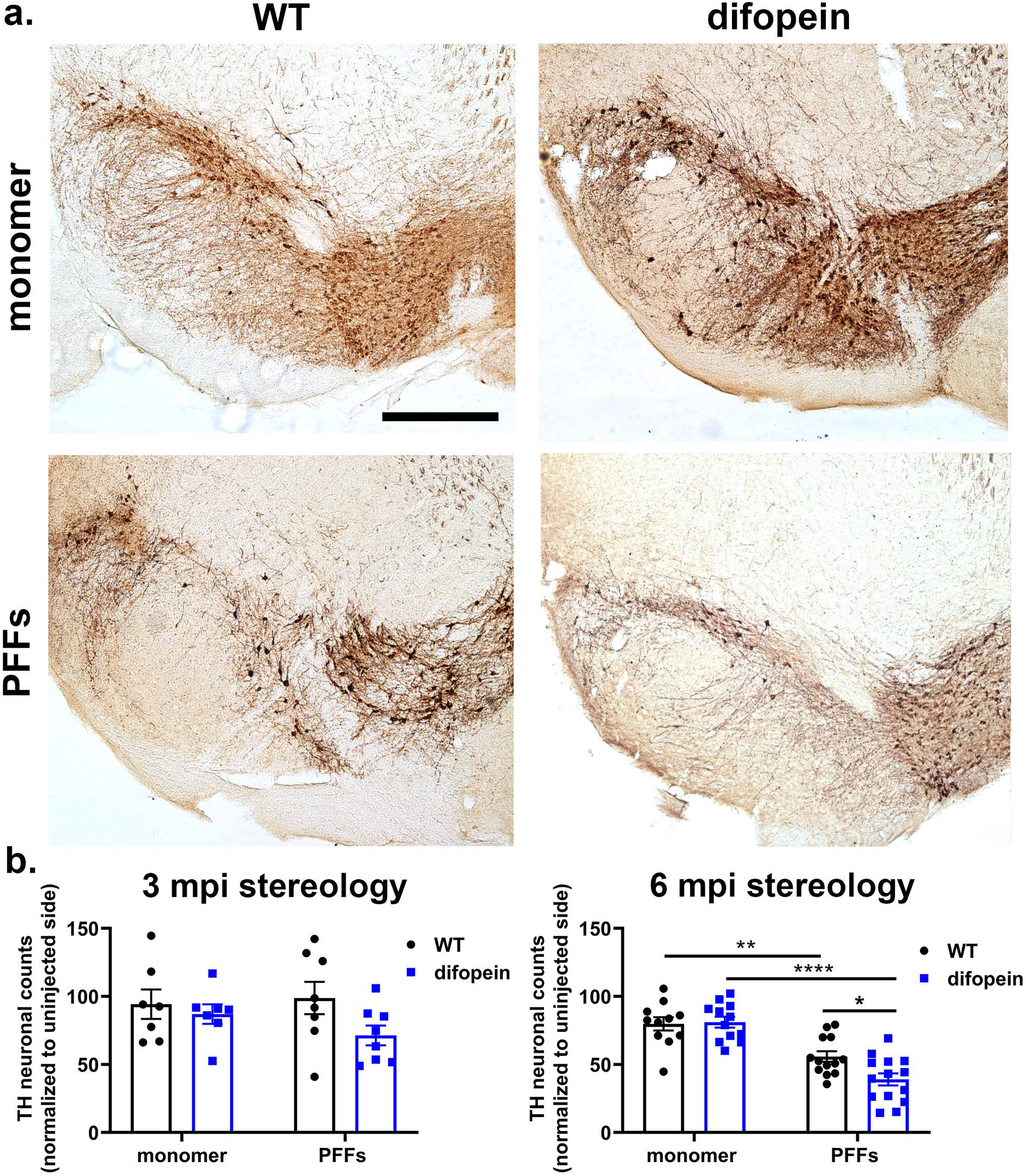
14-3-3 inhibition exacerbates nigral dopaminergic loss in response to PFFs. a) Representative images of TH immunohistochemistry in the SN of WT and difopein nigral mice (line 166) injected with αsyn monomers or PFFs at 6 mpi. Scale bar = 500 μm. b) Difopein expression in the nigra increases dopaminergic cell loss in response to PFFs. Stereological counts of TH-positive neurons in the SN of WT and difopein mice injected with αsyn monomers or PFFs at 3 mpi and 6 mpi. n=7-8 per group at 3 mpi; n=11-14 at 6 mpi. *p<0.05, **p<0.01, ****p<0.0001 (Tukey’s multiple comparison test). Error bars represent SEM.

To assess neuronal loss in the cortex of WT and transgenic difopein mice, we measured counts of layer IV pyramidal neurons. Since inclusion formation primarily occurs in layer IV and V pyramidal neurons [40], we used NECAB1 as a marker for layer IV pyramidal neurons and estimated neuronal death by counts of NECAB1-positive neurons per area (Supp. Fig. 5b). At 6 mpi, we observed a non-significant reduction in NECAB1-positive neuronal density in WT mice injected with PFFs compared to those injected with monomeric αsyn, while difopein mice injected with PFFs did show a significant reduction in NECAB1-positive neuronal density compared to difopein mice injected with monomeric αsyn (2-way ANOVA: genotype F (1, 27) = 1.426, p = 0.2428; PFF treatment F (1, 27) = 35.68, p < 0.0001; interaction F (1, 27) = 4.681, p = 0.0395; Supp. Fig. 5a, c). While not statistically significant, there was a slight trend towards decreased NECAB1 counts in difopein mice injected with PFFs compared to WT mice injected with PFFs (Supp. Fig. 5a, c). These findings in the SN and cortex suggest that the reduction in αsyn inclusion counts in difopein-expressing mice at 6 mpi is likely due to the increased loss of neurons in difopein mice compared to WT mice.

## Discussion

In this study we examined the role of 14-3-3s on αsyn aggregation and neuronal toxicity in the *in vivo* pre-formed fibril (PFF) mouse model. We tested the effects of 14-3-3θ overexpression and 14-3-3 inhibition by the pan-peptide inhibitor difopein in either the cortex or the SN at 2 time points after PFF injection. 14-3-3θ overexpression in the cortex and amygdala, as demonstrated in our 14-3-3θ transgenic line, delayed αsyn inclusion formation and rescued the social dominance deficit. 14-3-3θ overexpression in the SN by AAV also delayed αsyn inclusion formation and partially rescued dopaminergic neuronal loss. In contrast, difopein expression accelerated αsyn inclusion formation, accelerated the social dominance behavioral deficit, and increased neuronal loss. These results suggest that 14-3-3s act as a chaperone to reduce αsyn aggregation and its resulting toxicity on behavioral function and neuron loss.

We and other groups have demonstrated that inclusion formation in the mouse PFF model peaks around 3 months after PFF injection with a subsequent reduction in inclusions at later time points [1, 18]. This reduction in aggregates coincides with the onset of neuronal loss at 6 mpi [1, 18], suggesting that the reduction in aggregate numbers is due to the death of neurons with inclusions. Indeed, live imaging has demonstrated that neurons expressing αsyn inclusions ultimately die when tracked over time [25]. In our study, we found that 14-3-3θ overexpression decreased αsyn aggregation at 3 mpi, but interestingly increased αsyn aggregate counts at 6 mpi in areas expressing higher levels of 14-3-3θ. Conversely, 14-3-3 inhibition increased αsyn aggregation at 3 mpi but decreased it at 6 mpi in areas expressing difopein. We hypothesize that the subsequent increase in inclusion numbers with 14-3-3θ overexpression at 6 mpi is due to a delay in neuronal loss as compared to that seen in WT mice. Conversely, the decrease in inclusion numbers in difopein mice at 6 mpi is secondary to an acceleration of cell death relative to WT mice. Indeed, our analyses of neuronal loss and behavior are consistent with this interpretation. 14-3-3θ overexpression limited dopaminergic neuron loss in the SN at 6 mpi, and behavioral rescue also points to rescue of neuronal function. Conversely, 14-3-3 inhibition increased dopaminergic neuron loss at 6 mpi. Although not significant, 14-3-3 inhibition trended towards increasing dopaminergic neuron loss at 3 mpi as well, suggesting an acceleration of neuronal cell death. NECAB1-positive cell counts in the sensorimotor cortex were also reduced in PFF-injected difopein mice. The acceleration of the social dominance defect in difopein mice is consistent with this acceleration of neuron loss.

In order to assess the effects of 14-3-3s on αsyn toxicity in the PFF model, we used both transgenic and AAV-induced expression mouse models. Reproducible and consistent findings between AAV and transgene methods of 14-3-3θ overexpression suggest that our findings are not an artifact of transgene expression. The use of both methods also allowed for the assessment of selective areas of expression in modulating the effects of αsyn after initial aggregation initiated by PFF injection in the striatum, as αsyn inclusion formation is seen throughout multiple brain regions in both PD patients and in the PFF model. Our findings demonstrate that 14-3-3s’ effects on αsyn are not restricted to particular brain regions, but that 14-3-3s can impact αsyn pathology in multiple areas in which αsyn pathology is observed in the PFF model. Of note, we tested the effects of the overexpression of a single isoform 14-3-3θ in comparison to pan inhibition of 14-3-3 isoforms by difopein. Our lab has previously established the integral role of 14-3-3θ in modulating αsyn spread and toxicity [46], although other isoforms may also impact αsyn pathology. For example, 14-3-3η can regulate αsyn aggregation *in vitro* [28]. We used the pan-14-3-3 peptide inhibitor to eliminate the possibility that other 14-3-3 isoforms could compensate for lack of a single isoform.

14-3-3 proteins interact with multiple aggregation-prone proteins in neurodegenerative diseases, including tau, huntingtin and αsyn [9, 11, 17, 34, 41, 44]. Our lab previously established that 14-3-3θ acts as a chaperone to regulate αsyn seeding, cell-to-cell transmission, and toxicity in both the paracrine αsyn and *in vitro* PFF models [46]. 14-3-3θ complexes with αsyn to prevent its adoption of a pathologic conformation, limiting further αsyn aggregation. 14-3-3θ-complexed αsyn decreases its uptake, seeding potential, and paracrine toxicity. Here we further confirm the essential role for 14-3-3θ in the *in vivo* PFF model and predict that its protective effects *in vivo* involve its role as a chaperone. 14-3-3s could also act as a chaperone to regulate other aggregation-prone proteins, yet whether 14-3-3s do regulate other aggregation-prone proteins *in vivo* is yet to be determined. *In vitro* studies evaluating the impact of 14-3-3s on tau and huntingtin aggregation have been mixed [2, 24, 26, 30, 32, 47].

Based on our findings here and in other studies, we propose that disruption of 14-3-3 function may serve to promote the neurodegenerative process in PD and DLB. Alterations in 14-3-3s have been noted in PD models and in human disease [6, 20, 35, 48, 49]. We have previously shown that increased αsyn levels reduces 14-3-3θ expression in αsyn cellular and mouse models and that 14-3-3 levels are reduced in DLB [6, 20, 48, 49]. Additionally, we have observed increased 14-3-3θ phosphorylation in PD models and in human PD and DLB brains [20, 35]. A reduction in 14-3-3 levels and aberrant 14-3-3 phosphorylation may impair the chaperone function of 14-3-3θ to minimize αsyn misfolding. Future studies are in progress to examine the impact of 14-3-3 phosphorylation on αsyn pathology.

Motor phenotypes in the PFF model have been variably reported, with some groups finding strong motor deficits and others observing none [12, 18, 40]. Our data showed no consistent deficits in PFF-injected mice by wire hang, pole test, or rotarod at 6 mpi. Variability in behavioral deficits may be due to differences in protocol or in the genetic background of the mice used in each study. This lack of replication may also be due to differences in the synthesis of injected fibrils, resulting in different rates of αsyn aggregation and neuronal cell death. It has been previously established that αsyn aggregates reach different peak times in each brain region and then decrease as the aggregates are cleared and cells die [1, 18, 25, 27]. As a result, behavioral phenotypes in this model may vary based on the seeding potential of the PFFs injected. Interestingly, we consistently observed deficits in the social dominance tube test in PFF-injected mice at 6 mpi, pointing to strong implications for prefrontal cortical and amygdala involvement in this model, reflecting a disease profile reminiscent of DLB [40].

## Conclusion

In conclusion, we found that 14-3-3θ overexpression reduced behavioral deficits, delayed αsyn aggregation, and partially reduced neuronal loss, while 14-3-3 inhibition accelerated behavioral deficits, αsyn aggregation, and neuronal loss in the PFF mouse model. Our work here further demonstrates the neuroprotective effects of 14-3-3θ overexpression in multiple brain regions, indicating that this protective mechanism applies broadly to multiple cell types affected by αsyn pathology. Together these data indicate the role of 14-3-3s in the regulation of αsyn pathology and their therapeutic potential as a molecular target for synucleinopathies. Induction of 14-3-3θ may prove to be a viable technique for slowing disease progression.

## List of abbreviations

AAV: adenovirus associated virus
αsyn: alpha-synuclein
CTX: cortex
DLB: Dementia with Lewy Bodies
IHC: immunohistochemistry
LB: Lewy body
mpi: months post injection
PD: Parkinson’s disease
PFF: pre-formed fibrils
pS129-: phosphoserine 129
SN: substantia nigra
TH: tyrosine hydroxylase

## Declarations

### Ethics approval and consent to participate

Mice were used in accordance with the guidelines of the National Institute of Health (NIH) and University of Alabama at Birmingham (UAB) Institutional Animal Care and Use Committee (IACUC) and all experiments abided by the principles outlined in the Basel Declaration.

### Consent for publication

Not applicable

### Availability of data and material

The datasets generated and analyzed during the current study are available from the corresponding author on reasonable request.

### Authors’ contributions

RU designed and performed experiments, analyzed data, and wrote manuscript. MG designed and performed experiments, analyzed data, and reviewed manuscript. NK, AK, SC, and AP performed experiments and reviewed manuscript. TY designed experiments, analyzed data, and wrote and finalized manuscript. All authors read and approved the final manuscript.

## Acknowledgements

We are grateful to Drs. Laura Volpicelli-Daley and Andrew West for provision of recombinant αsyn. We thank Dr. Yi Zhou at Florida State University who provided the difopein mouse lines that were used in these studies.

## SUPPLEMENTAL FIGURES

**Supplemental Figure 1.**
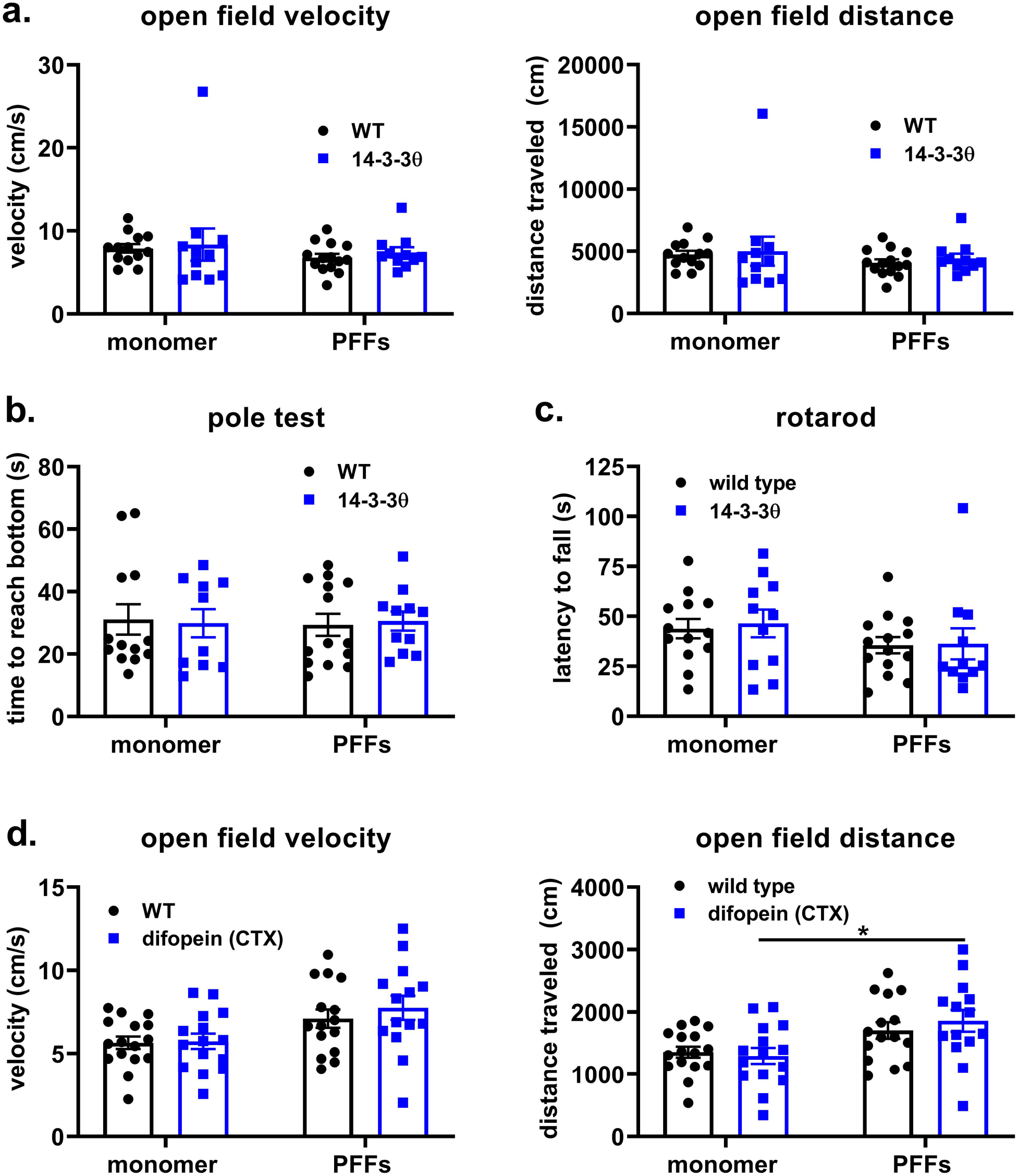
Motor behaviors are not affected by PFF treatment in transgenic 14-3-3θ or difopein mice. a) Quantification of average velocity and distance traveled in the open field test for WT and 14-3-3θ mice injected with αsyn monomer or PFFs at 6 mpi. n=11-14 per group, n.s. (Tukey’s multiple comparison test). Error bars represent SEM. b) Quantification of time to reach the bottom in the pole test for WT and 14-3-3θ mice injected with αsyn monomer or PFFs at 6 mpi. n=11-14 per group, n.s. (Tukey’s multiple comparison test). Error bars represent SEM. c) Quantification of latency to fall in the accelerating rotarod test on the second assessment day after 3 days of training for WT and 14-3-3θ mice injected with αsyn monomer or PFFs at 6 mpi. n=11-14 per group, n.s. (Tukey’s multiple comparison test). Error bars represent SEM. d) Quantification of average velocity and distance traveled in the open field test for WT and difopein (“cortical” line 138) mice injected with αsyn monomer or PFFs at 6 mpi. n=14-16 per group, *p<0.05 (Tukey’s multiple comparison test). Error bars represent SEM.

**Supplemental Figure 2.**
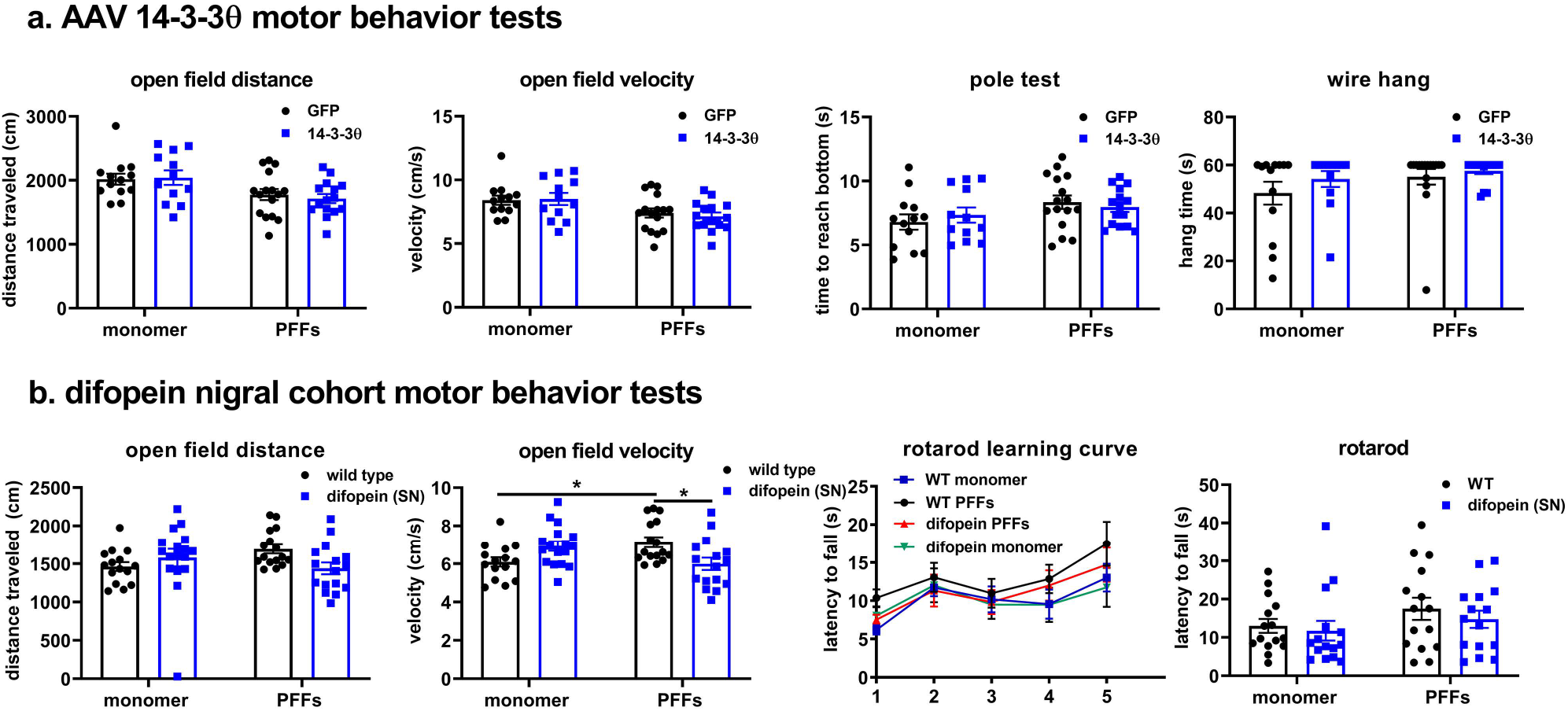
Motor behaviors are not affected by PFF treatment nor by 14-3-3 manipulation in the nigra. a) Quantification of average velocity and distance traveled in the open field test, time to reach the bottom in the pole test, and latency to fall on the wire hang test in AAV-GFP and AAV-14-3-3θ/GFP mice injected with αsyn monomer or PFFs at 6 mpi. n=12-16 per group. n.s. (Tukey’s multiple comparison test). Error bars represent SEM. b) Quantification of average velocity and distance traveled in the open field test and latency to fall in the accelerating rotarod test in WT and difopein (“nigral” line 166) mice injected with αsyn monomer or PFFs at 6 mpi. n=15-17 per group. *p<0.05 (Tukey’s multiple comparison test). Error bars represent SEM.

**Supplemental Figure 3.**
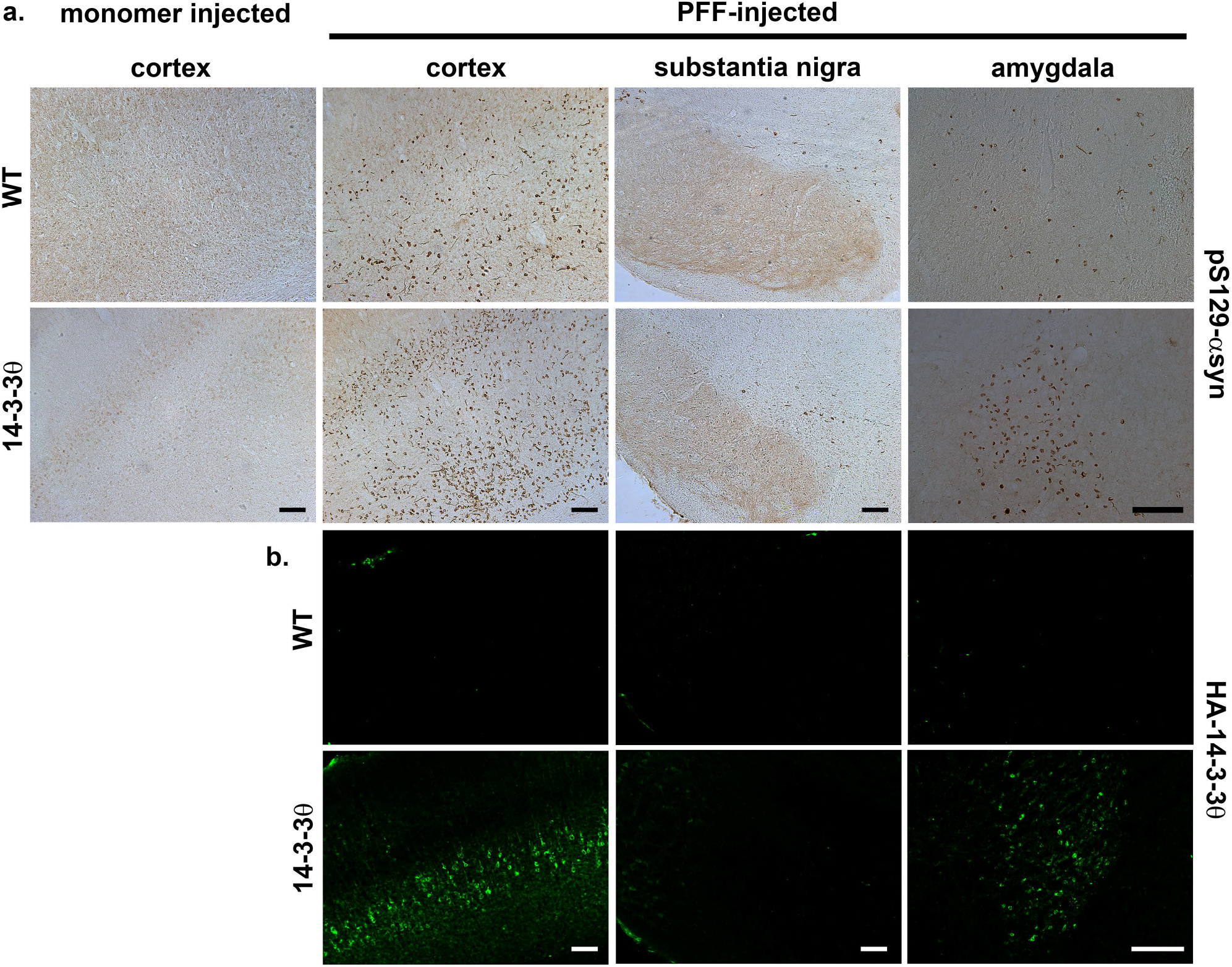
αSyn aggregation in 14-3-3θ transgenic mice at 6 months post injection. a) Representative images of pS129-αsyn immunostaining in WT and 14-3-3θ mice injected with αsyn monomer or PFFs in the cortex, SN, and amygdala at 6 mpi. Scale bar = 100 μm. b) Representative images of HA immunostaining in the cortex, SN, and amygdala demonstrates that HA-tagged 14-3-3θ is expressed in cortical and amygdala regions, but not in the SN. Scale bar = 100 μm.

**Supplemental Figure 4.**
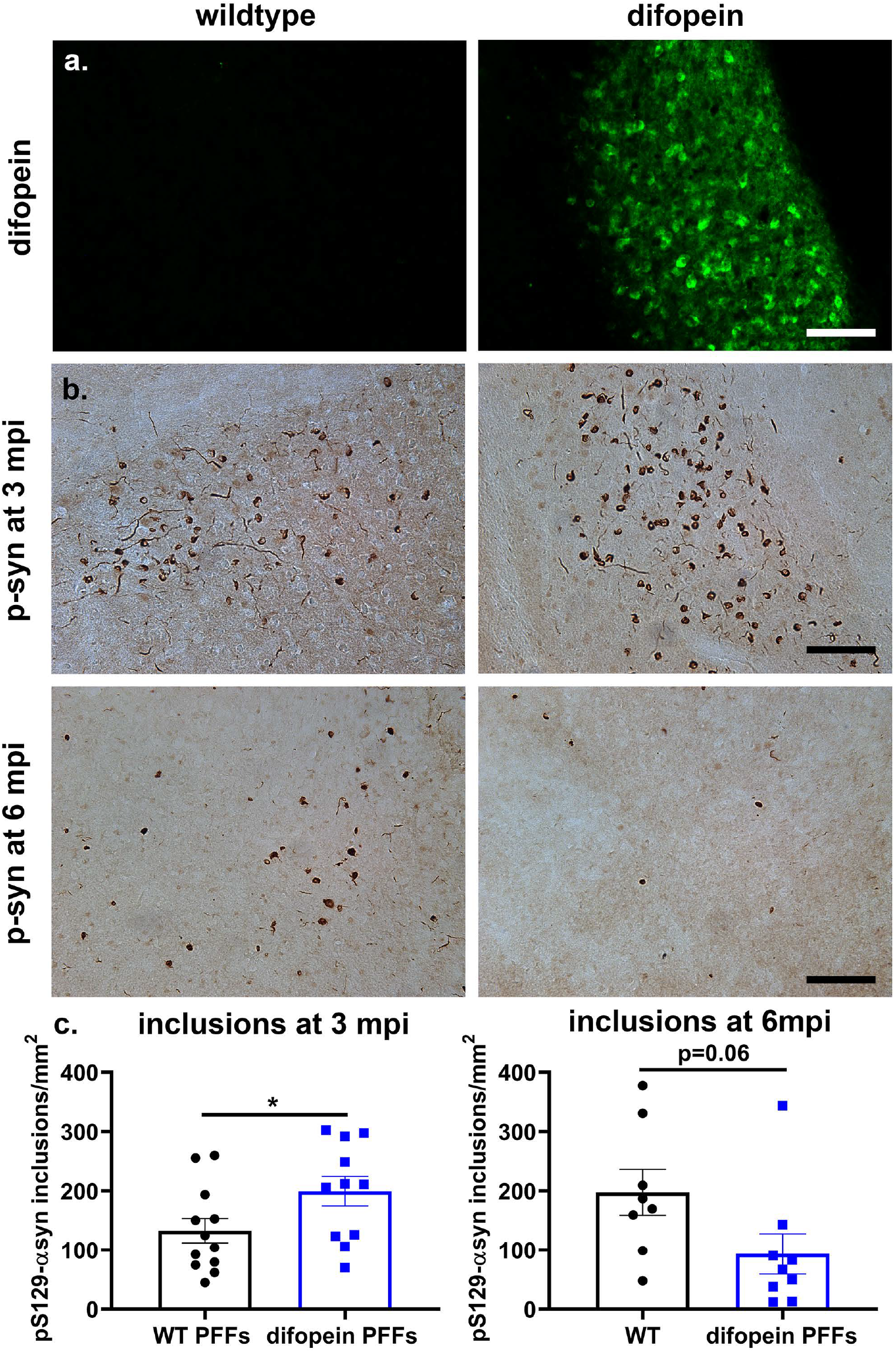
αSyn aggregation is accelerated in the amygdala in mice expressing difopein in the amygdala. a) Representative images of eYFP-difopein immunostaining in the amygdala of WT and difopein (“cortical” line 138) mice at 3 mpi. GFP-difopein expression is found only in difopein mice. Scale bar = 100 μm. b) Representative images of pS129-αsyn immunostaining in the cortex of WT and difopein mice at 3 and 6 mpi. Scale bar = 100 μm. c) Difopein expression increases inclusion counts in PFF-injected mice at 3 mpi. Quantification of pS129-αsyn positive inclusions at 3 mpi and 6 mpi in the amygdala of PFF-injected WT and difopein mice. n=11-12 per group at 3 mpi. n=8-9 at 6 mpi. *p<0.05 (Student’s t-test). Error bars represent SEM.

**Supplemental Figure 5.**
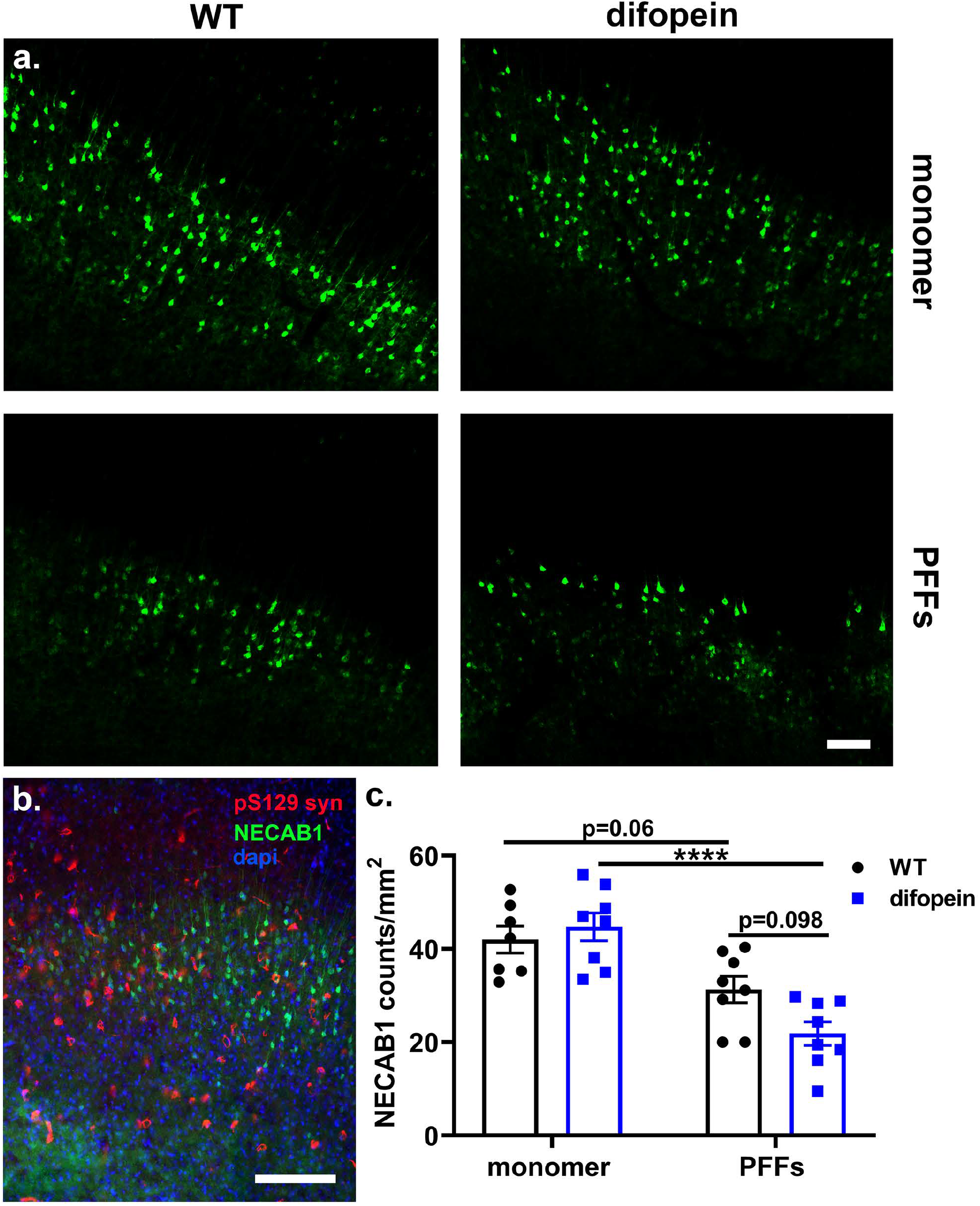
Difopein increases the loss of NECAB1-positive neurons in response to PFFs. a) Representative images of NECAB1-positive immunostaining in the cortex of WT and difopein mice injected with αsyn monomers or PFFs at 6 mpi. Scale bar = 100 μm. b) pS129-αsyn (red) immunostaining is concentrated in NECAB1-positive (green) IV and V layers of the sensorimotor cortex in a PFF-injected WT mouse. Scale bar = 100 μm. c) PFF-injected difopein mice have decreased NECAB1 counts in comparison to monomer-injected difopein mice. Quantification of NECAB1-positive neurons in the cortex of WT and difopein mice injected with αsyn monomers or PFFs at 6 mpi. n=7-8 per group. ****p<0.0001 (Tukey’s multiple comparison test). Error bars represent SEM.

## REFERENCES

1 Abdelmotilib H, Maltbie T, Delic V, Liu Z, Hu X, Fraser KB, Moehle MS, Stoyka L, Anabtawi N, Krendelchtchikova V (2017) α-Synuclein fibril-induced inclusion spread in rats and mice correlates with dopaminergic neurodegeneration. Neurobiology of disease 105: 84–98

2 Agarwal-Mawal A, Qureshi HY, Cafferty PW, Yuan Z, Han D, Lin R, Paudel HK (2003) 14-3-3 Connects Glycogen Synthase Kinase-3β to Tau within a Brain Microtubule-associated Tau Phosphorylation Complex. Journal of Biological Chemistry 278: 12722–12728 Doi 10.1074/jbc.M211491200

3 Arrant AE, Filiano AJ, Warmus BA, Hall AM, Roberson ED (2016) Progranulin haploinsufficiency causes biphasic social dominance abnormalities in the tube test. Genes, Brain and Behavior 15: 588–603

4 Berg D, Riess O, Bornemann A (2003) Specification of 14-3-3 proteins in Lewy bodies. Annals of neurology 54: 135–135

5 Cabin DE, Shimazu K, Murphy D, Cole NB, Gottschalk W, McIlwain KL, Orrison B, Chen A, Ellis CE, Paylor R (2002) Synaptic vesicle depletion correlates with attenuated synaptic responses to prolonged repetitive stimulation in mice lacking α-synuclein. Journal of Neuroscience 22: 8797–8807

6 Ding H, Fineberg NS, Gray M, Yacoubian TA (2013) α-Synuclein Overexpression Represses 14-3-3θ Transcription. Journal of Molecular Neuroscience 51: 1000–1009 Doi 10.1007/s12031-013-0086-5

7 Ding H, Underwood R, Lavalley N, Yacoubian TA (2015) 14-3-3 inhibition promotes dopaminergic neuron loss and 14-3-3θ overexpression promotes recovery in the MPTP mouse model of Parkinson’s disease. Neuroscience 307: 73–82

8 Emamzadeh FN (2016) Alpha-synuclein structure, functions, and interactions. J Res Med Sci 21: 29–29 Doi 10.4103/1735-1995.181989

9 Foote M, Zhou Y (2012) 14-3-3 proteins in neurological disorders. Int J Biochem Mol Biol 3: 152–164

10 Giasson BI, Murray IV, Trojanowski JQ, Lee VM-Y (2001) A hydrophobic stretch of 12 amino acid residues in the middle of α-synuclein is essential for filament assembly. Journal of Biological Chemistry 276: 2380–2386

11 Hashiguchi M, Sobue K, Paudel HK (2000) 14-3-3ζ Is an Effector of Tau Protein Phosphorylation. Journal of Biological Chemistry 275: 25247–25254 Doi 10.1074/jbc.M003738200

12 Hayakawa H, Nakatani R, Ikenaka K, Aguirre C, Choong C-J, Tsuda H, Nagano S, Koike M, Ikeuchi T, Hasegawa M et al (2020) Structurally Distinct α-Synuclein Fibrils Induce Robust Parkinsonian Pathology. Movement Disorders 35: 256–267 Doi 10.1002/mds.27887

13 Kajiwara Y, Buxbaum JD, Grice DE (2009) SLITRK1 binds 14-3-3 and regulates neurite outgrowth in a phosphorylation-dependent manner. Biological psychiatry 66: 918–925

14 Kawamoto Y, Akiguchi I, Nakamura S, Honjyo Y, Shibasaki H, Budka H (2002) 14-3-3 Proteins in Lewy Bodies in Parkinson Disease and Diffuse Lewy Body Disease Brains. Journal of Neuropathology & Experimental Neurology 61: 245–253 Doi 10.1093/jnen/61.3.245

15 Klein SM, Vykoukal J, Lechler P, Zeitler K, Gehmert S, Schreml S, Alt E, Bogdahn U, Prantl L (2012) Noninvasive in vivo assessment of muscle impairment in the mdx mouse model – A comparison of two common wire hanging methods with two different results. Journal of Neuroscience Methods 203: 292–297 Doi https://doi.org/10.1016/j.jneumeth.2011.10.001

16 Lavalley NJ, Slone SR, Ding H, West AB, Yacoubian TA (2016) 14-3-3 Proteins regulate mutant LRRK2 kinase activity and neurite shortening. Human molecular genetics 25: 109–122

17 Layfield R, Fergusson J, Aitken A, Lowe J, Landon M, Mayer RJ (1996) Neurofibrillary tangles of Alzheimer’s disease brains contain 14-3-3 proteins. Neuroscience Letters 209: 57–60 Doi https://doi.org/10.1016/0304-3940(96)12598-2

18 Luk KC, Kehm V, Carroll J, Zhang B, O’Brien P, Trojanowski JQ, Lee VM (2012) Pathological alpha-synuclein transmission initiates Parkinson-like neurodegeneration in nontransgenic mice. Science 338: 949–953 Doi 338/6109/949 [pii] 10.1126/science.1227157

19 Luk KC, Kehm V, Carroll J, Zhang B, O’Brien P, Trojanowski JQ, Lee VMY (2012) Pathological α-synuclein transmission initiates Parkinson-like neurodegeneration in nontransgenic mice. Science (New York, NY) 338: 949–953 Doi 10.1126/science.1227157

20 McFerrin MB, Chi X, Cutter G, Yacoubian TA (2017) Dysregulation of 14-3-3 proteins in neurodegenerative diseases with Lewy body or Alzheimer pathology. Annals of Clinical and Translational Neurology 4: 466–477 Doi 10.1002/acn3.421

21 Miczek KA, Brykczynski T, Grossman SP (1974) Differential effects of lesions in the amygdala, periamygdaloid cortex, and stria terminalis on aggressive behaviors in rats. J Comp Physiol Psychol 87: 760–771 Doi 10.1037/h0036971

22 Mrowiec T, Schwappach B (2006) 14-3-3 proteins in membrane protein transport. Biological chemistry 387: 1227–1236

23 Nemani VM, Lu W, Berge V, Nakamura K, Onoa B, Lee MK, Chaudhry FA, Nicoll RA, Edwards RH (2010) Increased Expression of α-Synuclein Reduces Neurotransmitter Release by Inhibiting Synaptic Vesicle Reclustering after Endocytosis. Neuron 65: 66–79 Doi https://doi.org/10.1016/j.neuron.2009.12.023

24 Omi K, Hachiya NS, Tanaka M, Tokunaga K, Kaneko K (2008) 14-3-3zeta is indispensable for aggregate formation of polyglutamine-expanded huntingtin protein. Neuroscience Letters 431: 45–50 Doi https://doi.org/10.1016/j.neulet.2007.11.018

25 Osterberg Valerie R, Spinelli Kateri J, Weston Leah J, Luk Kelvin C, Woltjer Randall L, Unni Vivek K (2015) Progressive Aggregation of Alpha-Synuclein and Selective Degeneration of Lewy Inclusion-Bearing Neurons in a Mouse Model of Parkinsonism. Cell Reports 10: 1252–1260 Doi https://doi.org/10.1016/j.celrep.2015.01.060

26 Papanikolopoulou K, Grammenoudi S, Samiotaki M, Skoulakis EM (2018) Differential effects of 14-3-3 dimers on Tau phosphorylation, stability and toxicity in vivo. Human molecular genetics 27: 2244–2261

27 Patterson JR, Duffy MF, Kemp CJ, Howe JW, Collier TJ, Stoll AC, Miller KM, Patel P, Levine N, Moore DJ et al (2019) Time course and magnitude of alpha-synuclein inclusion formation and nigrostriatal degeneration in the rat model of synucleinopathy triggered by intrastriatal α-synuclein preformed fibrils. Neurobiology of Disease 130: 104525 Doi https://doi.org/10.1016/j.nbd.2019.104525

28 Plotegher N, Kumar D, Tessari I, Brucale M, Munari F, Tosatto L, Belluzzi E, Greggio E, Bisaglia M, Capaldi S et al (2014) The chaperone-like protein 14-3-3η interacts with human α-synuclein aggregation intermediates rerouting the amyloidogenic pathway and reducing α-synuclein cellular toxicity. Human Molecular Genetics 23: 5615–5629 Doi 10.1093/hmg/ddu275

29 Qiao H, Foote M, Graham K, Wu Y, Zhou Y (2014) 14-3-3 Proteins Are Required for Hippocampal Long-Term Potentiation and Associative Learning and Memory. The Journal of Neuroscience 34: 4801–4808 Doi 10.1523/jneurosci.4393-13.2014

30 Qureshi HY, Li T, MacDonald R, Cho CM, Leclerc N, Paudel HK (2013) Interaction of 14-3-3ζ with microtubule-associated protein tau within Alzheimer’s disease neurofibrillary tangles. Biochemistry 52: 6445–6455

31 Ramser EM, Buck F, Schachner M, Tilling T (2010) Binding of αII spectrin to 14-3-3β is involved in NCAM-dependent neurite outgrowth. Molecular and Cellular Neuroscience 45: 66–74

32 Sadik G, Tanaka T, Kato K, Yamamori H, Nessa BN, Morihara T, Takeda M (2009) Phosphorylation of tau at Ser214 mediates its interaction with 14-3-3 protein: implications for the mechanism of tau aggregation. Journal of Neurochemistry 108: 33–43 Doi 10.1111/j.1471-4159.2008.05716.x

33 Shandala T, Woodcock JM, Ng Y, Biggs L, Skoulakis EM, Brooks DA, Lopez AF (2011) Drosophila 14-3-3∊ has a crucial role in anti-microbial peptide secretion and innate immunity. Journal of cell science 124: 2165–2174

34 Shirasaki Dyna I, Greiner Erin R, Al-Ramahi I, Gray M, Boontheung P, Geschwind Daniel H, Botas J, Coppola G, Horvath S, Loo Joseph A et al (2012) Network Organization of the Huntingtin Proteomic Interactome in Mammalian Brain. Neuron 75: 41–57 Doi https://doi.org/10.1016/j.neuron.2012.05.024

35 Slone SR, Lavalley N, McFerrin M, Wang B, Yacoubian TA (2015) Increased 14-3-3 phosphorylation observed in Parkinson’s disease reduces neuroprotective potential of 14-3-3 proteins. Neurobiology of Disease 79: 1–13 Doi https://doi.org/10.1016/j.nbd.2015.02.032

36 Slone SR, Lesort M, Yacoubian TA (2011) 14-3-3theta protects against neurotoxicity in a cellular Parkinson’s disease model through inhibition of the apoptotic factor Bax. PLoS One 6: e21720

37 Soumiya H, Godai A, Araiso H, Mori S, Furukawa S, Fukumitsu H (2016) Neonatal Whisker Trimming Impairs Fear/Anxiety-Related Emotional Systems of the Amygdala and Social Behaviors in Adult Mice. PloS one 11: e0158583–e0158583 Doi 10.1371/journal.pone.0158583

38 St Martin JL, Klucken J, Outeiro TF, Nguyen P, Keller-McGandy C, Cantuti-Castelvetri I, Grammatopoulos TN, Standaert DG, Hyman BT, McLean PJ (2007) Dopaminergic neuron loss and up-regulation of chaperone protein mRNA induced by targeted over-expression of alpha-synuclein in mouse substantia nigra. Journal of neurochemistry 100: 1449–1457

39 Steidinger TU, Slone SR, Ding H, Standaert DG, Yacoubian TA (2013) Angiogenin in Parkinson Disease Models: Role of Akt Phosphorylation and Evaluation of AAV-Mediated Angiogenin Expression in MPTP Treated Mice. PLOS ONE 8: e56092 Doi 10.1371/journal.pone.0056092

40 Stoyka LE, Arrant AE, Thrasher DR, Russell DL, Freire J, Mahoney CL, Narayanan A, Dib AG, Standaert DG, Volpicelli-Daley LA (2020) Behavioral defects associated with amygdala and cortical dysfunction in mice with seeded α-synuclein inclusions. Neurobiology of disease 134: 104708

41 Umahara T, Uchihara T, Tsuchiya K, Nakamura A, Iwamoto T, Ikeda K, Takasaki M (2004) 14-3-3 proteins and zeta isoform containing neurofibrillary tangles in patients with Alzheimer’s disease. Acta neuropathologica 108: 279–286

42 Vincenz C, Dixit VM (1996) 14-3-3 proteins associate with A20 in an isoform-specific manner and function both as chaperone and adapter molecules. Journal of Biological Chemistry 271: 20029–20034

43 Volpicelli-Daley LA, Luk KC, Lee VM (2014) Addition of exogenous α-synuclein preformed fibrils to primary neuronal cultures to seed recruitment of endogenous α-synuclein to Lewy body and Lewy neurite-like aggregates. Nat Protoc 9: 2135–2146 Doi 10.1038/nprot.2014.143

44 Waelter S, Boeddrich A, Lurz R, Scherzinger E, Lueder G, Lehrach H, Wanker EE (2001) Accumulation of mutant huntingtin fragments in aggresome-like inclusion bodies as a result of insufficient protein degradation. Molecular biology of the cell 12: 1393–1407

45 Wallén-Mackenzie Å, Nordenankar K, Fejgin K, Lagerström MC, Emilsson L, Fredriksson R, Wass C, Andersson D, Egecioglu E, Andersson M et al (2009) Restricted Cortical and Amygdaloid Removal of Vesicular Glutamate Transporter 2 in Preadolescent Mice Impacts Dopaminergic Activity and Neuronal Circuitry of Higher Brain Function. The Journal of Neuroscience 29: 2238–2251 Doi 10.1523/jneurosci.5851-08.2009

46 Wang B, Underwood R, Kamath A, Britain C, McFerrin MB, McLean PJ, Volpicelli-Daley LA, Whitaker RH, Placzek WJ, Becker K et al (2018) 14-3-3 Proteins Reduce Cell-to-Cell Transfer and Propagation of Pathogenic α-Synuclein. The Journal of Neuroscience 38: 8211–8232 Doi 10.1523/jneurosci.1134-18.2018

47 Xu Z, Graham K, Foote M, Liang F, Rizkallah R, Hurt M, Wang Y, Wu Y, Zhou Y (2013) 14-3-3 protein targets misfolded chaperone-associated proteins to aggresomes. Journal of Cell Science 126: 4173–4186 Doi 10.1242/jcs.126102

48 Yacoubian TA, Cantuti-Castelvetri I, Bouzou B, Asteris G, McLean PJ, Hyman BT, Standaert DG (2008) Transcriptional dysregulation in a transgenic model of Parkinson disease. Neurobiology of Disease 29: 515–528 Doi https://doi.org/10.1016/j.nbd.2007.11.008

49 Yacoubian TA, Slone SR, Harrington AJ, Hamamichi S, Schieltz JM, Caldwell KA, Caldwell GA, Standaert DG (2010) Differential neuroprotective effects of 14-3-3 proteins in models of Parkinson’s disease. Cell death & disease 1: e2–e2

50 Yano M, Nakamuta S, Wu X, Okumura Y, Kido H (2006) A novel function of 14-3-3 protein: 14-3-3ζ is a heat-shock–related molecular chaperone that dissolves thermal-aggregated proteins. Molecular biology of the cell 17: 4769–4779

51 Zhou T, Sandi C, Hu H (2018) Advances in understanding neural mechanisms of social dominance. Current Opinion in Neurobiology 49: 99–107 Doi https://doi.org/10.1016/j.conb.2018.01.006

